# Arabidopsis uORF-containing mRNAs behave differently from NMD targets

**DOI:** 10.1101/2021.09.16.460672

**Authors:** Hsin-Yen Larry Wu, Polly Yingshan Hsu

## Abstract

Upstream ORFs (uORFs) are common regulatory elements in the 5’ untranslated regions of eukaryotic mRNAs. In addition to repressing main ORF translation, uORF translation in animals also reduces mRNA stability through nonsense-mediated decay (NMD). In contrast, the roles of uORFs in plants are less understood. Here, we identified actively translated uORFs (TuORFs) in Arabidopsis through ribosome profiling and systematically examined their roles in gene expression. Like animal systems, Arabidopsis TuORFs are associated with 38%, 14%, and 43% reductions in translation efficiency, mRNA half-lives, and protein levels, respectively. However, we found TuORF-containing mRNAs have 51% higher transcript levels, and this phenomenon is persistent in diverse tissues and developmental stages across plants. We present multiple lines of evidence that indicate Arabidopsis uORF-containing mRNAs generally exhibit distinct behavior from known NMD targets. First, TuORF-containing mRNAs are not increased in NMD mutants. Second, TuORF-containing mRNAs and known NMD targets have distinct expression patterns, and they are translationally repressed via different mechanisms. Finally, TuORF- containing mRNAs and NMD targets are degraded through separate pathways. Our results suggest that Arabidopsis TuORFs reduce mRNA stability and translation through mechanisms different from NMD and highlight a fundamental difference in gene regulation mediated by TuORFs in plants and animals.

## INTRODUCTION

Upstream open reading frames (uORFs) are cis-acting translational repressors that regulate many eukaryotic protein-coding transcripts. Mutations or modifications of uORFs are associated with human diseases and plant traits (Calvo et al., 2009; Xu et al., 2017; Zhang et al., 2018; Lee et al., 2021). Considering AUG as a translation start codon, 30%–70% of genes contain at least one potential uORF in moss, Arabidopsis, fly, mouse, and human (Kim et al., 2007; Johnstone et al., 2016; Zhang et al., 2021). Genes with predicted uORFs are enriched for signal transduction proteins, transcription factors, and membrane proteins in animals (Zhang et al., 2021) and for transcription factors and protein kinases in plants (Kim et al., 2007), underlining the potential importance of uORFs in modulating the expression of vital cellular regulators.

Many studies in eukaryotes have reported that uORF translation reduces main ORF (mORF) translation (Chew et al., 2016; Johnstone et al., 2016; Liu et al., 2013; Wu et al., 2019; Lin et al., 2019), and it is estimated to reduce protein production by 30%–80% (Calvo et al., 2009; Chew et al., 2016). According to the scanning model of mRNA translation, the 43S pre-initiation complex, which contains the small ribosomal subunit, several translation initiation factors, and initiator tRNA, binds the 5’ cap and scans the 5’ untranslated region (UTR) to search for an optimal initiation sequence (Hinnebusch et al., 2016). When a uORF is translated, the ribosome may stall during elongation or at termination, or it may dissociate from the mRNA after termination, thus reducing the translation of the downstream mORF (Dever et al., 2020).

The mORFs downstream of uORFs may get translated through leaky scanning or reinitiation (Kozak, 1999).

uORF translation is also expected to trigger mRNA degradation, presumably through nonsense-mediated decay (NMD). NMD is an evolutionarily conserved regulatory mechanism to eliminate aberrant mRNAs, such as those that contain premature termination codons or unusually long 3’ UTRs (He and Jacobson, 2015; Kurosaki et al., 2019). Two pathways have been established in mammals: the exon junction complex (EJC)-dependent and long 3’ UTR pathways. Normally, EJC and two NMD factors (UP-FRAMESHIFT SUPPRESSOR 2 and 3; UPF2 and UPF3) deposited during splicing of mRNAs will be evicted by the ribosome in the first round of translation. In the presence of a premature termination codon, EJC-UPF2-UPF3 complexes downstream of the termination codon remain attached to the mRNA. This allows UPF1, which is part of the surveillance complex recruited at translation termination, to interact with UPF2 and UPF3 in proximity. The interaction of UPF1, UPF2, and UPF3 activates the SUPPRESSOR WITH MORPHOLOGICAL EFFECT ON GENITALIA1 (SMG1) kinase, which phosphorylates UPF1. This establishes the first phase of NMD, substrate recognition, and activates downstream signaling and eventually the second phase of NMD, substrate degradation, which involves SMG5, SMG6, and SMG7. Phosphorylated UPF1 also inhibits translation initiation (Isken et al., 2008). Alternatively, unusually long 3’ UTRs, which result in poor translation termination, are believed to promote NMD through phosphorylated UPF1 independent of EJC (Kurosaki et al., 2014).

The plant NMD pathway is best characterized in *Arabidopsis thaliana*. Arabidopsis possesses homologous NMD factors involved in the first phase, including UPF1, UPF2, and UPF3, and in the second phase, SMG7 (Raxwal and Riha, 2016). However, Arabidopsis is missing the homologs of animal SMG1, SMG5, and SMG6 (Lloyd and Davies, 2013). The absence of key NMD components in plants suggests significant variations in the NMD mechanisms between plants and animals. It remains an open question how the differences in NMD components affect substrate recognition and degradation in plants.

In Arabidopsis, putative NMD targets have been identified based on transcripts that overly accumulate in NMD-defective mutants (Kurihara et al., 2009; Kalyna et al., 2012; Drechsel et al., 2013; Degtiar et al., 2015). However, there is a concern that the upregulation of some transcripts in NMD mutants might be caused by secondary effects of NMD (Raxwal et al., 2020). For example, NMD mutants affect numerous aspects of plant physiology, and some alleles cause severe growth retardation and lethality due to constitutive immune responses (Jeong et al., 2011; Rayson et al., 2012; Riehs-Kearnan et al., 2012; Filichkin et al., 2015; Merchante et al., 2015; Chiam et al., 2019; Yoine et al., 2006). Moreover, UPF1 has been reported to have multiple functions in plants, such as alternative splicing regulation and translational control, in addition to NMD (Raxwal et al., 2020). To identify *bona fide* NMD targets, *upf1* and *smg7*, which are defective in the first and second phases of NMD, respectively, were introduced into *pad4*, in which the immune response is inactivated (Riehs-Kearnan et al., 2012; Raxwal et al., 2020). The 333 transcripts upregulated in both *upf1 pad4* and *smg7 pad4* were defined as high-confidence NMD targets. It was shown these NMD targets are translationally regulated by UPF1 and that their turnover requires decapping and 5’ to 3’ decay (Raxwal et al., 2020).

As uORF translation could mimic a transcript with an unusually long 3’ UTR, uORF-containing mRNAs are predicted to be targeted by NMD (Kertész et al., 2006; Kurosaki et al., 2019). Consistent with this expectation, genome-wide studies in zebrafish, mouse, and human have found that transcripts with translated uORFs have lower mRNA levels globally (Chew et al., 2016; Johnstone et al., 2016). Moreover, transcripts containing translated uORFs are upregulated in knockdown of NMD factors in mouse embryonic stem cells and human cell lines (Hurt et al., 2013; Baird et al., 2018), supporting the idea that these transcripts are NMD targets.

In plants, whether uORF translation triggers NMD is still under debate. Two genome-wide studies suggest that uORF-containing mRNAs are targets of NMD, as transcripts possessing predicted uORFs are enriched in the genes upregulated in Arabidopsis *upf1-1* and *upf3-1* mutants (Kurihara et al., 2009; Drechsel et al., 2013). In contrast, three studies suggested that NMD only controls a small fraction of uORF-containing mRNAs in plants, such as those containing uORFs with > 35 codons (∼2% of uORF-containing genes), or those responsive to certain environmental stimuli, such as polyamine or blue light (Nyikó et al., 2009; Uchiyama-Kadokura et al., 2014; Kurihara et al., 2018). Moreover, evidence from several gene-specific or genome-wide studies suggests that uORFs do not trigger NMD. First, the mRNA levels of several uORF-containing transcripts, including *NIP5;1* and *AtMHX*, are not increased in NMD mutants (Saul et al., 2009; Tanaka et al., 2016). Second, a genome-wide study reported that genes with increasing numbers of uORFs did not show lower transcript abundances in Arabidopsis and tomato (Li and Liu, 2020). Third, genes with predicted uORFs are not over- represented in the consensus NMD target list identified from three NMD mutants, including *upf1-5*, *upf3-1*, and *smg7-1* (Rayson et al., 2012). It was also reported that 47 mRNAs that contain Conserved Peptide uORFs (CPuORFs) behave differently from NMD targets prior to their degradation mediated by XRN4 (Nagarajan et al., 2019). It is unknown whether some uORF-containing mRNAs escape NMD or whether uORF-containing mRNAs are generally not targets of NMD in plants. It has been concluded that the mRNA features studied so far are insufficient to infer NMD targetability in plants (Ohtani and Wachter, 2019). Overall, it remains unclear how translated uORFs affect mRNA stability and what the role of NMD in this process is in plants.

One limitation of the previous studies is that predicted uORFs, which include both translated and untranslated uORFs, were commonly included in the analyses. Although many uORFs have been predicted based on nucleotide sequences (Zhang et al., 2021), determining which uORFs are actively translated still requires experimental evidence. Recently, deep sequencing of ribosome footprints – known as ribosome profiling or Ribo-seq – has been exploited to study genome-wide mRNA translation (Ingolia et al., 2009; Brar and Weissman, 2015). Several Ribo-seq studies in Arabidopsis have deployed different strategies to identify translated uORFs (TuORFs) (Liu et al., 2013; Juntawong et al., 2014; Hsu et al., 2016; Hu et al., 2016; Bazin et al., 2017; Kurihara et al., 2020). Among the parameters used, 3-nucleotide (3-nt) periodicity, which corresponds to translating ribosomes deciphering 3 nts per codon, is considered the most stringent criterion to identify high-confidence translation events (Ingolia et al., 2009; Calviello et al., 2016; Xu et al., 2018; Harnett et al., 2021). Our previous work and a recent study using RiboTaper (Calviello et al., 2016), which computes the statistical significance of 3-nt periodicity, uncovered 187–1378 AUG-initiated TuORFs in Arabidopsis (Hsu et al., 2016; Kurihara et al., 2020).

Here we further improved the Ribo-seq methodology and identified 2093 AUG-initiated TuORFs in Arabidopsis seedlings. We systematically examined the assumed roles of TuORFs in mORF translation and mRNA decay by integrating multiple genome-wide datasets. As predicted, we found that TuORF mRNAs are associated with lower translation efficiency, shorter mRNA half-lives, and lower protein abundance. Surprisingly, we found that, in contrast to the animal paradigm, plant TuORF mRNAs are associated with higher steady-state mRNA levels, and they are not increased in the NMD mutants. Examinations of additional parameters revealed that uORFs and NMD regulate gene expression through separate mechanisms. Taken together, our results reveal unexpected features and regulation of uORFs in Arabidopsis and suggest that plants and animals have evolved different uORF-dependent mechanisms for gene regulation.

## RESULTS

### Translated uORFs identified via an improved Ribo-seq method

Previously we optimized Ribo-seq to reach high 3-nt periodicity, and we exploited this unique feature of mRNA translation to identify translated ORFs in Arabidopsis (Hsu et al., 2016). Although the data revealed many unannotated translation events, the number of TuORFs uncovered was low, presumably due to insufficient Ribo-seq coverage within uORFs. We proposed that increasing Ribo-seq coverage could improve the efficiency of translated ORF identification, especially for relatively short ORFs, such as uORFs. Here we optimized the coverage of Ribo-seq in 7-day-old Arabidopsis seedlings by minimizing the potential loss of ribosome footprints during the purification process (see METHODS for details).

In addition to high correlations among biological replicates (**Figure S1A**), our new Ribo-seq data displayed characteristics expected for high-quality data, including strong 3-nt periodicity (92% in-frame reads), high enrichment for coding sequences (CDSs), and characteristic ribosome footprint lengths (**Figure 1A-C**). The improved coverage is evident when examining the number of unique ribosome footprints detected globally. At a sequencing depth of 20 million reads from randomly selected reads, the number of unique ribosome footprints in our new data is 4.01-fold higher than that in our previous dataset (**Figure 1D**). The coverage improvement is also evident in individual transcript profiles within both uORF and mORF regions. For example, a TuORF in *REPRESSOR OF GA1* (*RGA1*) was identified in our current data but missed in our previous study despite the mRNA levels being similar (**Figure S1B**).

**Figure 1.**
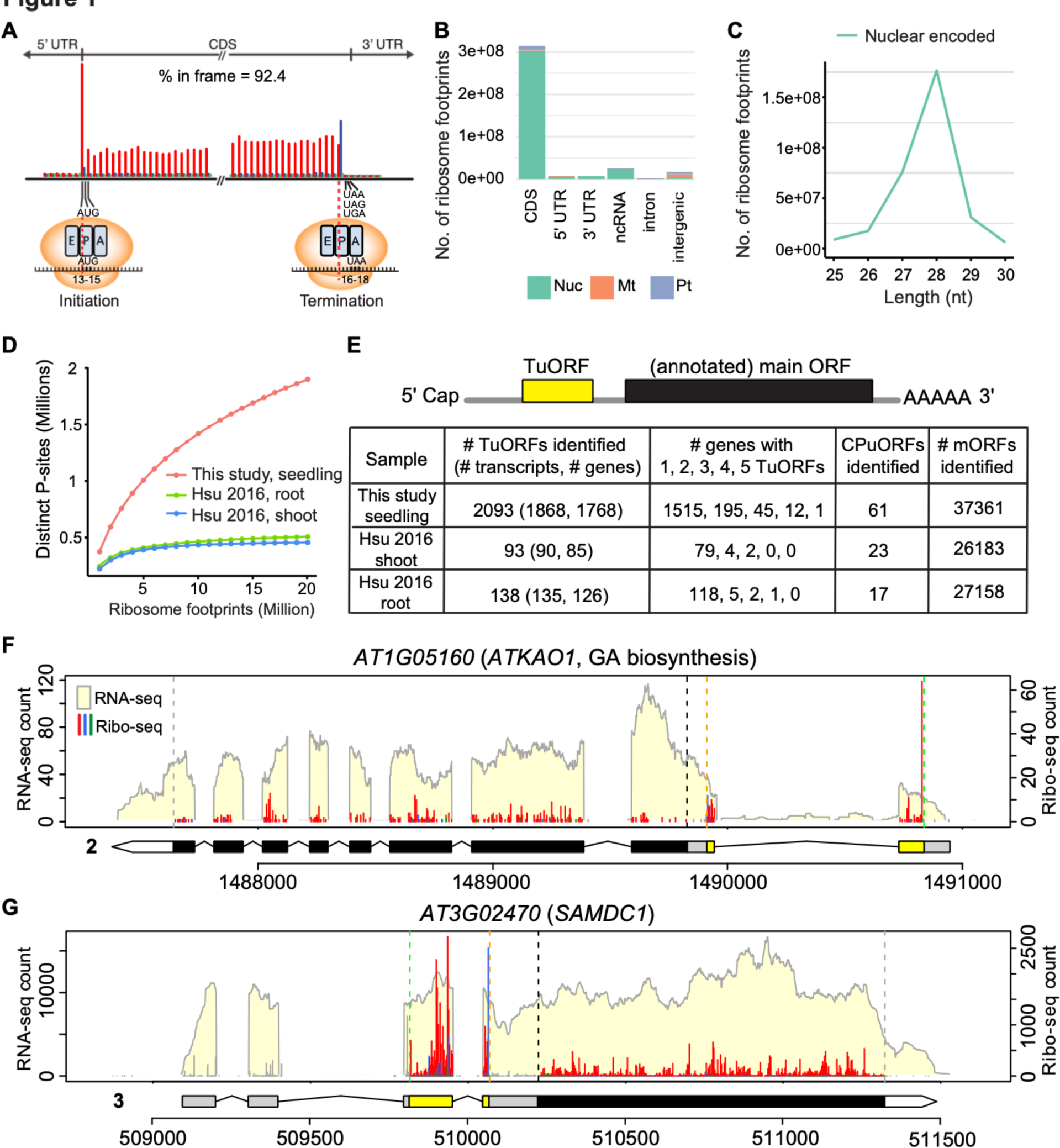
Enhanced Ribo-seq coverage improves uORF identification. (A) Metagene analysis of 28-nt ribosome footprints mapped to regions near the start and stop codons of annotated ORFs. The reads are presented with their first nt at the P-site, which is the 13th nt for 28-nt footprints. The reads are colored in red, blue, and green to indicate they are in the first (expected), second, and third reading frames, respectively. The majority of footprints were mapped to the CDS in the expected reading frame (92.4% in frame). (B) Genomic features mapped by ribosome footprints. Reads that mapped to nuclear (Nuc)-, mitochondrial (Mt)-, and plastid (Pt)-encoded genes are shown. (C) Length distribution of ribosome footprints. Reads that mapped to nuclear-encoded genes are presented. (D) Distinct P-sites detected in 1 to 20 million randomly selected ribosome footprints from our current and previous datasets. (E) Numbers of TuORFs, CPuORFs, and mORFs identified in our current and previous datasets. (F-G) RNA-seq and Ribo-seq profiles of *ATKAO1 and SAMDC1* from our current data. RNA-seq coverage is shown with a light-yellow background. Ribo-seq reads are presented with their first nt at the P-site, and they are colored in red, blue, and green to indicate they are in the first (expected), second, and third reading frames, respectively. Within the gene models, yellow boxes represent the TuORFs, black boxes represent the annotated mORFs, and gray and white regions indicate 5’ UTRs and 3’ UTRs, respectively. The specific isoform being considered is indicated to the left of the gene model. Within the profiles, green and orange vertical dashed lines represent translation start and stop, respectively, for the TuORF. Black and gray vertical dashed lines represent translation start and stop, respectively, for the annotated mORF.

The high-coverage data increased the number of TuORFs uncovered. In total, 2093 TuORFs were identified in 1768 genes based on significant 3-nt periodicity using RiboTaper (with the default cutoff of RiboTaper, Multitaper F-test, p < 0.05) and Araport11 annotation (**Figure 1E** and **Table S1A**). This result is in strong contrast to the relatively few TuORFs identified in Arabidopsis seedling shoots (93 TuORFs) and roots (138 TuORFs) from our previous data obtained using plants grown under similar conditions and using the same statistical analysis and annotations (**Figure 1E** and **Table S1B-C**). The new data also revealed translation from a higher number of Conserved Peptide uORFs (CPuORFs) and annotated coding sequences (mORFs of protein-coding genes) (**Figure 1E** and **Table S1D-F**). We manually plotted all TuORF genes to confirm strong 3-nt periodicity present in the expected uORF regions using RiboPlotR (Wu and Hsu, 2021) (examples shown in **Figure 1F-G, Figure S2**). Taken together, these results show that our enhanced-coverage Ribo-seq data dramatically improved the identification of TuORFs.

Previous analyses of predicted uORFs found that transcription factors and protein kinases are overrepresented among uORF-containing genes (Von Arnim et al., 2014; Kim et al., 2007). Consistent with these analyses, functions related to transcription factors and protein kinases/phosphatases were identified as being enriched in our Gene Ontology (GO) analysis of TuORF-containing genes (**Table S2**). We found that many TuORF-containing genes encode extensively studied regulators of plant growth and environmental responses. For example, this list included well-known transcription factors, such as those involved in hormone signaling (RGA1 and AUXIN RESPONSE FACTOR2/3/8), root development (SCARECROW), the circadian clock (TIMING OF CAB EXPRESSION1), and light signaling (PHYTOCHROME INTERACTING FACTOR3/4/7) (Silverstone et al., 1998; Strayer et al., 2000; Di Laurenzio et al., 1996; Ellis et al., 2005; Nagpal et al., 2005; Ni et al., 1998; Huq and Quail, 2002; Leivar et al., 2008). The list also included well-characterized protein kinases, such as Phytochromes B and E, ribosomal-protein S6 kinases (S6K1/2), and SNF1-related protein kinase 2 (SNRK2.1/2.5) (**Figure S1B, S2A-N**) (Sharrock and Quail, 1989; Devlin et al., 1998; Zhang et al., 1994; Mizoguchi et al., 1995; Park et al., 1993; Boudsocq et al., 2004). These results support the notion that TuORFs control important cellular regulators in plants.

### TuORF-containing genes have higher mRNA abundance but are associated with lower translation efficiency, shorter mRNA half-lives, and lower protein abundance

To understand how uORF translation affects gene expression, we compared mRNA abundance and translation efficiency of TuORF-containing genes (hereafter, TuORF genes), genes containing sequence-predicted uORFs but <u>u</u>ndetected by RiboTaper (hereafter, UuORF genes), and genes containing an annotated 5’ UTR but no potential uORFs (hereafter, no-uORF genes). Note that some UuORFs may be translated but missed in our analysis because their 3- nt periodicity is insufficient to pass the statistical test: these include extremely short uORFs, those translated at a low frequency, and those overlapping with mORF or other uORFs and thus disrupting the 3-nt periodicity. To study the global effects of uORFs on mRNA stability and protein abundance, we also compared genes with TuORFs, UuORFs, or no-uORFs, using available mRNA half-life data determined by metabolic labeling in Arabidopsis seedlings and two quantitative proteomics datasets from shoots and roots of Arabidopsis seedlings (Szabo et al., 2020; Song et al., 2018). The growth conditions and source of the data are summarized in **Table S3**. Below we presented the data in cumulative plots to show the trends of different gene groups and in box plots to show the statistical results (**Figure 2A-H and S3**).

**Figure 2.**
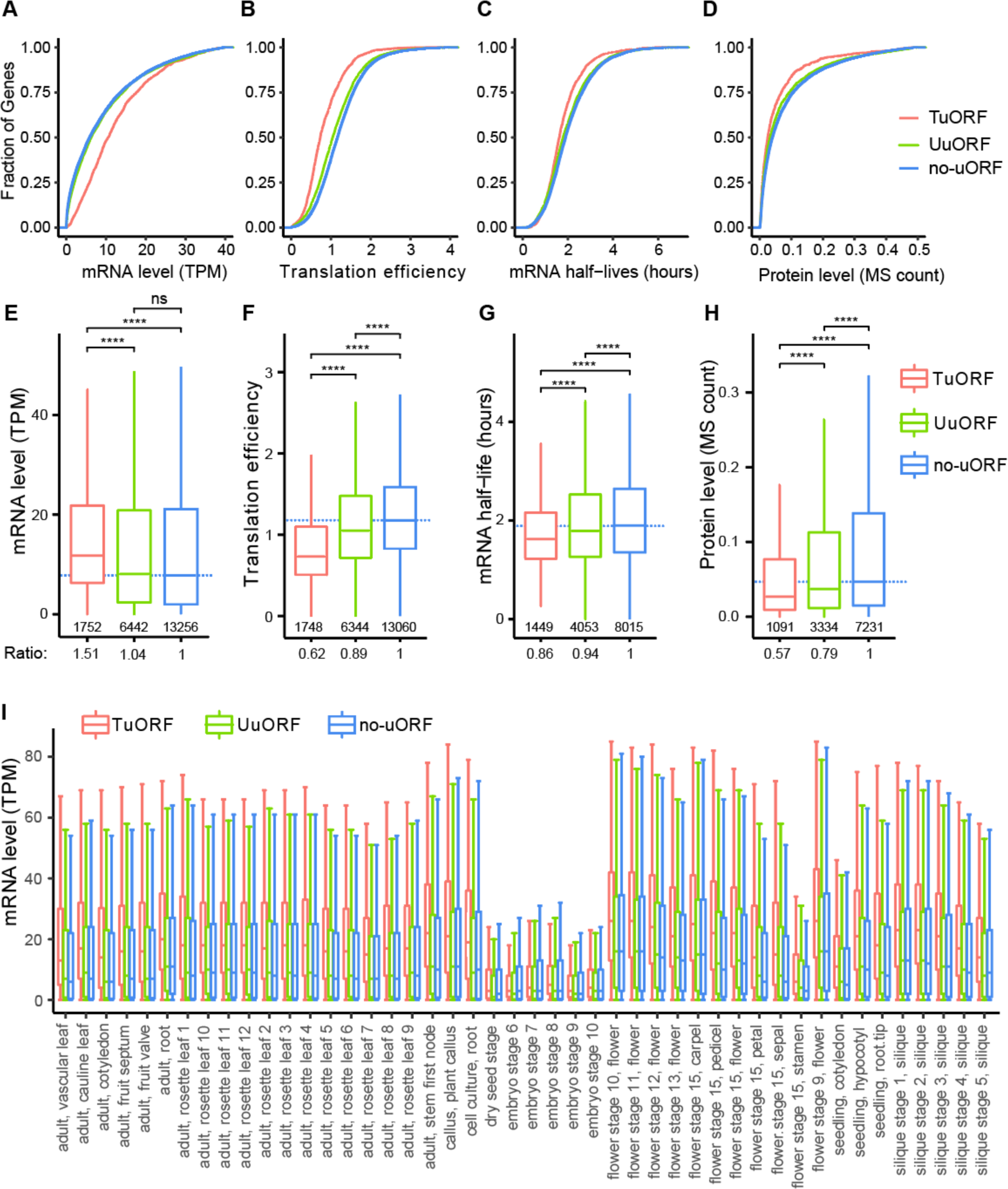
TuORF-containing genes have higher mRNA abundance but are associated with lower translation efficiency, shorter mRNA half-lives, and lower protein abundance. (A-D) Cumulative plots and (E-H) boxplots for the mRNA levels, translation efficiency, mRNA half-lives, and protein levels of TuORF, UuORF, and no-uORF genes in Arabidopsis seedlings. The mRNA half-life data, which were measured with metabolic labeling in Arabidopsis seedlings, were extracted from (Szabo et al., 2020), and the protein abundance as determined by quantitative proteomics in Arabidopsis seedling shoots were extracted from (Song et al., 2018). Within the boxplots (E-H), the dashed lines mark the median level of the no-uORF genes; the number of genes in each category is listed below the lower whisker. The ratios underneath the plots indicate the median of each group normalized to that of the no-uORF genes. The statistical significance was determined by Wilcoxon signed-rank test. The adjusted p-value was determined using the Benjamini and Yekutieli procedure to control for the false discovery rate in multiple testing (* 0.05 > p > 0.01, ** 0.01 > p > 0.001, *** 0.001 > p > 1e-4, **** 1e-4 > p > 0). These statistical analyses and graphic layouts were applied to all boxplots in the study. (I) The mRNA levels of TuORF, UuORF, and no-uORF genes in 46 Arabidopsis tissues and developmental stages from independent studies. The TPM values from RNA-seq quantification were downloaded from the EMBO-EBI expression atlas (Papatheodorou et al., 2020). Additional 82 Arabidopsis and tomato RNA-seq datasets were presented in Figure S5.

Studies in vertebrates have shown that uORF translation is associated with lower steady-state mRNA abundance, consistent with the expectation that TuORF mRNAs are regulated by NMD (Chew et al., 2016; Johnstone et al., 2016). Surprisingly, we found that TuORF genes in Arabidopsis are associated with higher steady-state mRNA levels than those of UuORF and no-uORF genes (**Figure 2A, 2E**), despite having shorter mRNA half-lives as expected (**Figure 2C, 2G**). The median mRNA level of TuORF genes is 51% higher than that of no-uORF genes (**Figure 2E**). The higher steady-state mRNA levels and shorter mRNA half-lives together suggest that TuORF genes are highly transcribed. To examine whether the high transcript levels of TuORF genes are conserved in other plant species, we reanalyzed our previous tomato Ribo-seq data from seedling roots (Wu et al., 2019) and found the TuORF genes in tomato also have higher steady-state mRNA levels relative to no-uORF genes (**Figure S4A**). Given that Arabidopsis and tomato diverged more than 100 million years ago (Clarke et al., 2011), our results suggest that TuORF-associated higher mRNA abundance could be prevalent in plants. To investigate whether our observation is specific to seedlings or common to different developmental stages/tissues/genetic backgrounds in Arabidopsis and tomato, we examined the mRNA levels of the uORF-containing genes in 128 independent RNA-seq studies collected by the EMBO-EBI expression atlas (Papatheodorou et al., 2020). Consistent with our Arabidopsis seedlings and tomato seedling root data, we found the TuORF genes are associated with higher mRNA levels in diverse tissues, developmental stages, and ecotypes in Arabidopsis and tomato (**Figure 2I and S5A-C**), supporting the hypothesis that TuORF genes are preferentially expressed at high levels in these plants under a wide range of sampling conditions.

CPuORFs are uORFs that encode evolutionarily conserved peptides, and they were identified independently from our approach by evolutionary conservation (Hayden and Jorgensen, 2007; Jorgensen and Dorantes-Acosta, 2012; Takahashi et al., 2012; Vaughn et al., 2012; Van Der Horst et al., 2019). Many CPuORFs were also identified as TuORFs in our study (**Figure 1E, 1G**). We found that the median mRNA abundance of CPuORF genes is 2.2-fold of that of no-uORF genes (**Figure S6**), consistent with our observations that genes with TuORFs are expressed at higher levels.

Next, we examined the translation efficiency, mRNA half-lives, and protein levels of TuORF, UuORF, and no-uORF genes. Consistent with the animal studies, we found Arabidopsis TuORF genes are associated with lower translation efficiency, shorter mRNA half-lives, and less protein accumulation than UuORF and no-uORF genes (**Figure 2B-D, 2F-H, S3, S4B**). Interestingly, even though globally TuORF genes start with 51% more mRNAs than no-uORF genes (**Figure 2E**), they produce 41%–43% less protein (**Figure 2H, S3**), on average, likely due to the combined effects of a 38% reduction in translation efficiency (**Figure 2F**) and a 14% reduction in mRNA half-lives (**Figure 2G**). These observations suggest that the global repression of TuORF gene protein synthesis is determined by multilayer post-transcriptional regulation. Notably, for the four parameters examined here, UuORF genes consistently have levels between those of TuORF genes and no-uORF genes: they show a 4% increase in mRNA levels and 11%, 6%, and 21% reductions in translation efficiency, mRNA half-lives, and protein abundance, respectively (**Figure 2A-H**). These results indicate that UuORFs have smaller effects on gene expression than TuORFs and highlight the importance of distinguishing TuORFs and UuORFs in genome-wide analysis.

### TuORF genes are larger and likely evolutionarily older than other genes

We next investigated whether there are general structural differences among TuORF, UuORF, and no-uORF mRNAs. Considering the directionality of ribosome scanning, we hypothesized TuORFs might be associated with longer 5’ UTRs. Indeed, we found that the median 5’ UTR length of TuORF mRNAs is 2.5-fold of that of no-uORF mRNAs (**Figure S7A**), which is consistent with a previous report that uORFs in longer 5’ UTRs are more likely to be translated (Liu et al., 2013). However, unexpectedly, we found that TuORF mRNAs also have longer CDSs and 3’UTRs, as well as longer transcript lengths, than UuORF mRNAs (**Figure S7B-D**). No-uORF mRNAs are the shortest in all categories examined (**Figure S7A-D**). These results support the hypothesis that TuORFs are more likely to be found in longer 5’ UTRs and reveal that TuORF mRNAs tend to encode larger proteins (**Figure S7B**). The enrichment of TuORF genes for transcription factors, protein kinases, and related functions in GO term analysis (**Table S2**) also implies TuORF genes tend to encode proteins with conserved domains. It is known that evolutionarily older genes are generally longer, expressed at higher levels, and have protein sequences that are under stronger purifying selection (Wolfa et al., 2009; Cui et al., 2015). Our observations that TuORF genes possess many features of evolutionarily older genes imply that older genes are more likely to be regulated through mRNA translation and degradation mediated by TuORFs.

### TuORF mRNAs are not upregulated in the NMD mutants

The established paradigm in vertebrates is that uORF translation triggers NMD and thus results in low steady-state mRNA levels (Chew et al., 2016; Johnstone et al., 2016; Hurt et al., 2013; Baird et al., 2018). Despite having higher steady-state mRNA levels (**Figure 2A and 2E**), TuORF genes in Arabidopsis also have shorter mRNA half-lives (**Figure 2C and 2G**). Therefore, we investigated whether TuORF-containing transcripts in Arabidopsis are still subject to NMD.

If TuORFs trigger NMD, one would expect TuORF mRNAs to be among the known NMD targets and their mRNA levels to be increased in NMD-deficient mutants. We first examined whether TuORF mRNAs were reported in the high-confidence NMD target list (Raxwal et al., 2020). Among the 333 NMD targets that were upregulated in both *upf1 pad4* and *smg7 pad4* adult leaves (Raxwal et al., 2020), 248 genes with an annotated 5’ UTR were expressed in our data (**Table S4A**). However, only 49 of 1748 TuORF genes were identified in the high-confidence NMD target list (**Table S4B**). The 49 genes represent 2.8% of TuORF-containing genes and 20% of the NMD targets. These results suggest that TuORFs generally do not trigger NMD; rather, NMD targets are more likely to be regulated by TuORFs than other genes (Chi-square test with Yates’ correction: p-value = 7.555e-10). One caveat is that the poor overlap between the two gene lists could be due to differences in growth conditions, developmental stages, and sample handling in different labs (**Table S3**).

To further investigate if TuORF mRNAs are NMD targets, we examined whether they are upregulated in NMD-deficient mutants by analyzing four published RNA-seq datasets for NMD mutants (Drechsel et al., 2013; Merchante et al., 2015; Raxwal et al., 2020; Gloggnitzer et al., 2014). Although these datasets were collected from different NMD mutants and at various developmental stages (**Table S3**), the high-confidence NMD targets are consistently and significantly upregulated in all the NMD mutants examined compared with the control plants (i.e., 3.05-, 1.54-, 4.98-, and 2.65-fold for NMD targets without TuORFs; 3.04-, 2.23-, 5.04-, and 2.66-fold for NMD targets with TuORFs) (**Figure 3A-D**). In contrast, TuORF mRNA levels show no or little change in the NMD mutants (i.e., 1.01-, 0.98-, 1.10- and 1.10-fold). Thus, only the NMD targets, but not TuORF mRNAs, are dramatically increased in the NMD mutants, supporting the idea that uORF translation in Arabidopsis may not trigger NMD.

**Figure 3:**
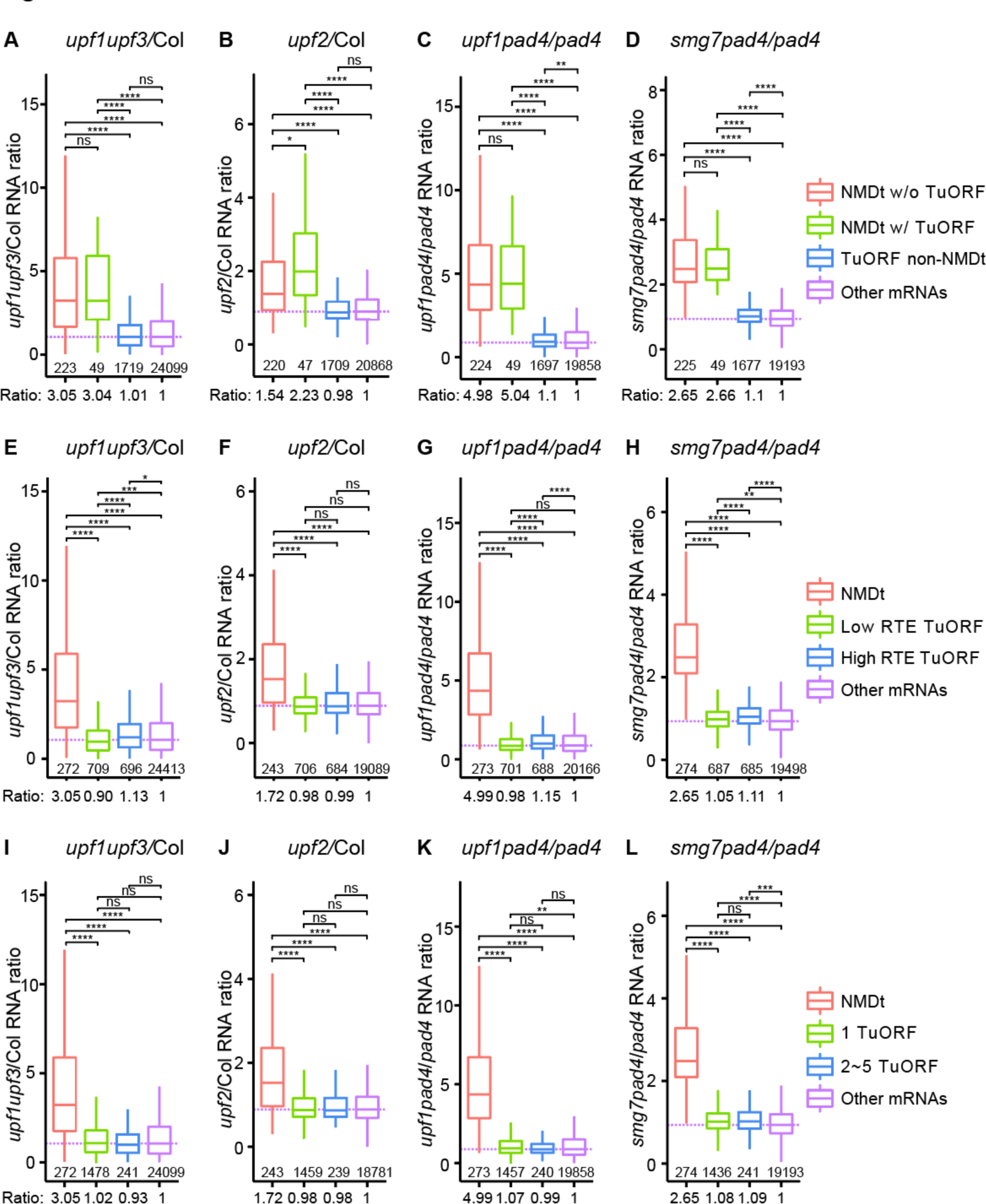
TuORF mRNAs are not upregulated in the NMD mutants. The ratios of mRNA levels between NMD mutants and their control plants from four RNA-seq datasets (Drechsel et al., 2013; Merchante et al., 2015; Raxwal et al., 2020; Gloggnitzer et al., 2014) were used to examine the effects of the NMD pathway on four gene groups: NMD targets without TuORF (NMDt w/o TuORF), NMD targets with TuORF (NMDt w/ TuORF), TuORF mRNAs not identified as NMD targets (TuORF non-NMDt) and other mRNAs. (A) *upf1* (*lba1* allele) *upf3-1* double mutant compared with wild type (Col-0). (B) *upf2-10* compared with wild type (Col-0). (C) *upf1-3 pad4* compared with *pad4*. (D) *smg7-1 pad4* compared with *pad4*. See Table S3 for specific growth conditions for each dataset. (E-H) Same datasets as 4A-D, but the TuORF mRNAs were grouped based on low or high relative TuORF translation efficiency (RTE) (i.e., TE TuORF / TE mORF). (I-L) Same datasets as 4A-D, but the TuORF mRNAs were grouped based on the number of TuORF(s). Within the boxplots, the dashed lines mark the median level of other mRNAs; the number of genes in each category is listed below the lower whisker. The ratios underneath the plots indicate the median of each group normalized to that of other mRNAs. The statistical significance was determined as described in the Figure 2 legend.

Given different uORFs are translated at various levels (See **Figure 1F-G, S1B, S2A-N** for examples), it is possible only the highly repressive TuORFs efficiently trigger NMD. To test this hypothesis, we split the TuORF genes into two groups based on their uORF relative translation efficiency (RTE) (i.e., TE of an uORF is normalized to TE of its mORF). However, even the higher TuORF RTE group shows no or little effects on the mRNA accumulation in the NMD mutants (i.e., 1.13-, 0.99-, 1.15- and 1.11-fold) (**Figure 3E-H**). We also tested whether having more TuORFs in a transcript efficiently triggers NMD by splitting the TuORF genes into two groups based on the number of TuORFs detected. We found that even transcripts with 2∼5 TuORFs do not lead to significant mRNA increase in the NMD mutants, either (i.e., 0.93-, 0.98-, 0.99- and 1.09-fold)(**Figure 3I-L**). Together, these results support the idea that NMD is not a major mechanism for controlling uORF-containing mRNAs.

### Lengths of the TuORF and 3’ UTR alone are insufficient to predict NMD targetability

A previous study suggested that uORFs that encode a peptide longer than 35 amino acids are likely to trigger NMD in Arabidopsis (Nyikó et al., 2009). To test whether TuORF length is associated with NMD targetability, we analyzed TuORF lengths in the NMD targets and non-NMD targets. We found that the NMD target TuORF genes tend to have longer uORF peptides than non-NMD target TuORF genes (median length 28 amino acids vs. 20.5 amino acids, p = 9.874e-07, Wilcoxon signed-rank test). However, the distributions of the uORF peptide lengths in NMD targets and non-NMD targets largely overlap (**Figure S8**), indicating that TuORF length alone is insufficient to predict whether a TuORF triggers NMD of a transcript.

We also examined whether the high-confidence NMD targets have longer 3’ UTRs, a feature that potentially triggers NMD. We found that the NMD targets do not have significantly longer 3’ UTRs, regardless of whether they contain TuORFs (**Figure S9**). Taken together, our results are consistent with the notion that no definite RNA features studied to date are sufficient to predict NMD targets in plants (Ohtani and Wachter, 2019).

### TuORF mRNAs and NMD targets have distinct expression patterns

To further characterize the roles of NMD and TuORFs in gene expression regulation, we compared the expression patterns of the NMD targets and TuORF mRNAs (**Figure 4**).

**Figure 4.**
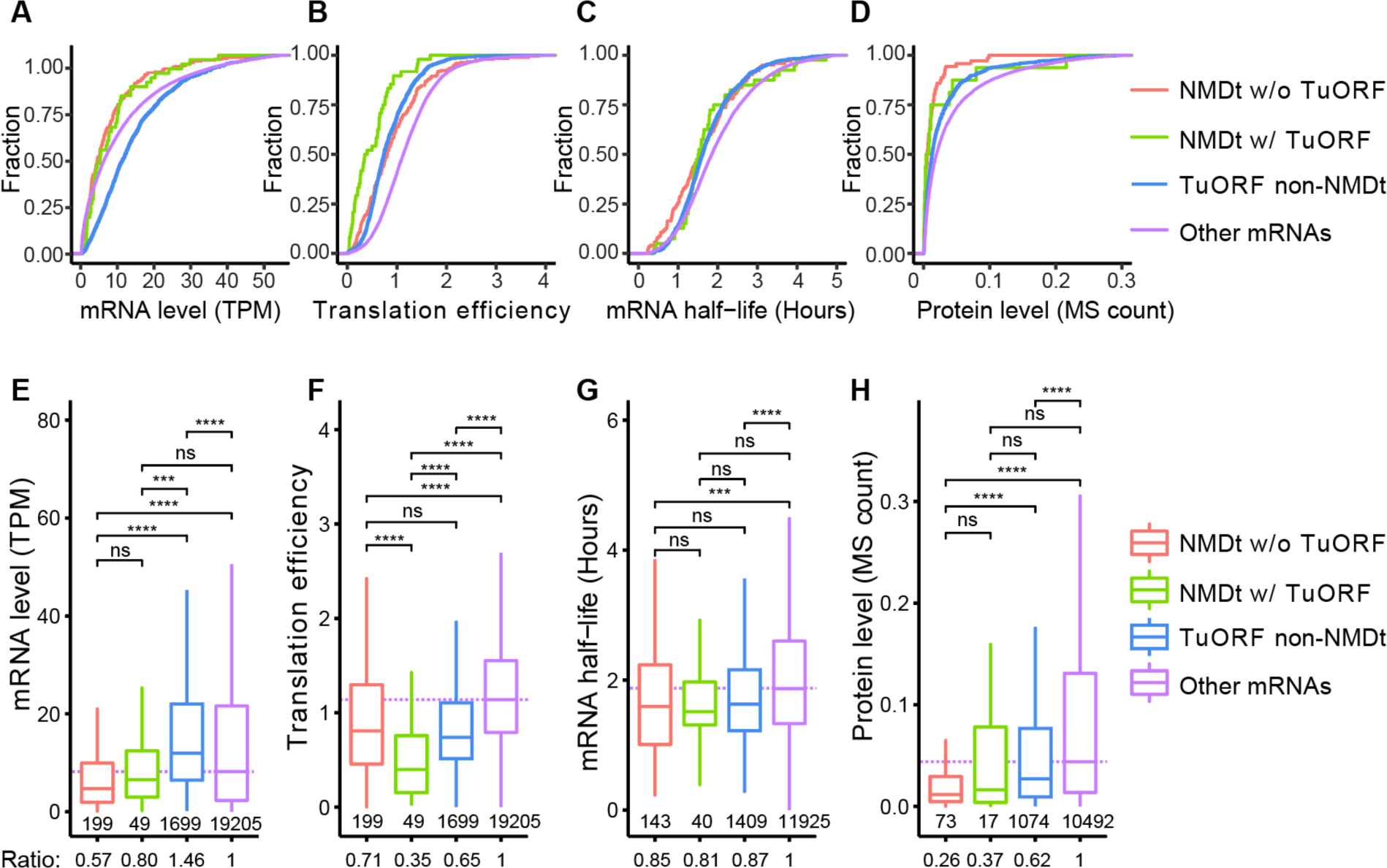
Expression patterns of NMD targets and TuORF mRNAs. Cumulative plots (A-D) and boxplots (E-H) for the mRNA levels (A, E), translation efficiency (B, F), mRNA half-lives (C, G), and protein levels (D, H) of NMD targets (NMDt) without or with TuORFs compared with those of TuORF mRNAs not identified as NMD targets and other mRNAs. Within the boxplots, the dashed lines mark the median level of other mRNAs; the number of genes in each category is listed below the lower whisker. The ratios underneath the plots indicate the median of each group normalized to that of other mRNAs. The statistical significance was determined as described in the Figure 2 legend.

Consistent with the expectation that NMD targets are being actively degraded, we found that NMD targets have lower mRNA abundance than other mRNAs; this is in strong contrast to TuORF mRNAs, which have higher mRNA abundance than other mRNAs (**Figure 4A, 4E**). While both NMD targets and TuORF mRNAs are associated with lower translation efficiency, shorter mRNA half-lives, and lower protein abundance than other mRNAs (**Figure 4B-D, 4F-H**), the NMD targets are associated with even lower protein levels than TuORF mRNAs (**Figure 4D, 4H**).

If TuORFs effectively trigger NMD, one would expect TuORF mRNAs to have translation efficiency similar to that of NMD targets with TuORFs, which would inherently be repressed by both TuORFs and NMD. However, we found that NMD targets with TuORFs have significantly lower translation efficiency than either NMD targets without TuORFs or TuORF mRNAs not identified as NMD targets (**Figure 4B, 4F**). This additive repression of translation efficiency in NMD targets with TuORFs suggests that NMD and TuORFs suppress translation through separate mechanisms. This result also supports that TuORFs do not effectively trigger NMD. Our observation is consistent with the expectation that TuORFs prevent the scanning pre-initiation complex from reaching the mORF, while NMD suppresses translation through UPF1 and/or other translational repressors (Isken et al., 2008; Raxwal et al., 2020; Zinshteyn et al., 2021).

Together, the differences in the expression patterns between TuORF mRNAs and NMD targets further support the idea that TuORF mRNAs behave differently from NMD targets in Arabidopsis.

### TuORF mRNAs and NMD targets are degraded through different mRNA decay mechanisms

We next investigated the mechanisms underlying the decay of TuORF mRNAs and NMD targets. mRNA decay can occur in the 5’ to 3’ or 3’ to 5’ direction, which is carried out by different machinery (Sieburth and Vincent, 2018). A genome-wide survey of mRNA decay rates in wild type and mutants defective in either 5’ to 3’ decay or 3’ to 5’ decay, or both, has been conducted in Arabidopsis (Sorenson et al., 2018). In that survey, the *vcs-7* mutant, a null mutant of *VARICOSE*, which encodes a scaffold protein for the mRNA decapping complex, was used to examine the 5’ to 3’ mRNA decay; the *sov* mutant, defective in *SUPPRESSOR OF VARICOSE*, which encodes a ribonuclease II, was used to examine 3’ to 5’ decay; and the *vcs/sov* double mutant was used to examine decay affected in both directions (Deyholos et al., 2003; Goeres et al., 2007; Xu et al., 2006; Zhang et al., 2010). We exploited this published dataset (Sorenson et al., 2018) to study the genetic requirements for the mRNA decay of TuORF mRNAs, NMD targets, TuORF-containing NMD targets, and other mRNAs.

We first compared the decay rates of each gene group across the four genetic backgrounds (**Figure 5**). Similar to mRNAs that are non-NMD targets and lack TuORFs (i.e., other mRNAs, **Figure 5D, 5H**), TuORF mRNAs have dramatically reduced decay rates in the *vcs* mutant compared to the wild type (**Figure 5A, 5E**), indicating that decapping and 5’ to 3’ decay are critical for degrading these transcripts. In the *vcs/sov* double mutant, TuORF mRNAs have slightly but significantly lower decay rates than in the *vcs* mutant (**Figure 5A and 5E**), suggesting *SOV* also plays a minor role in degrading TuORF mRNAs.

**Figure 5.**
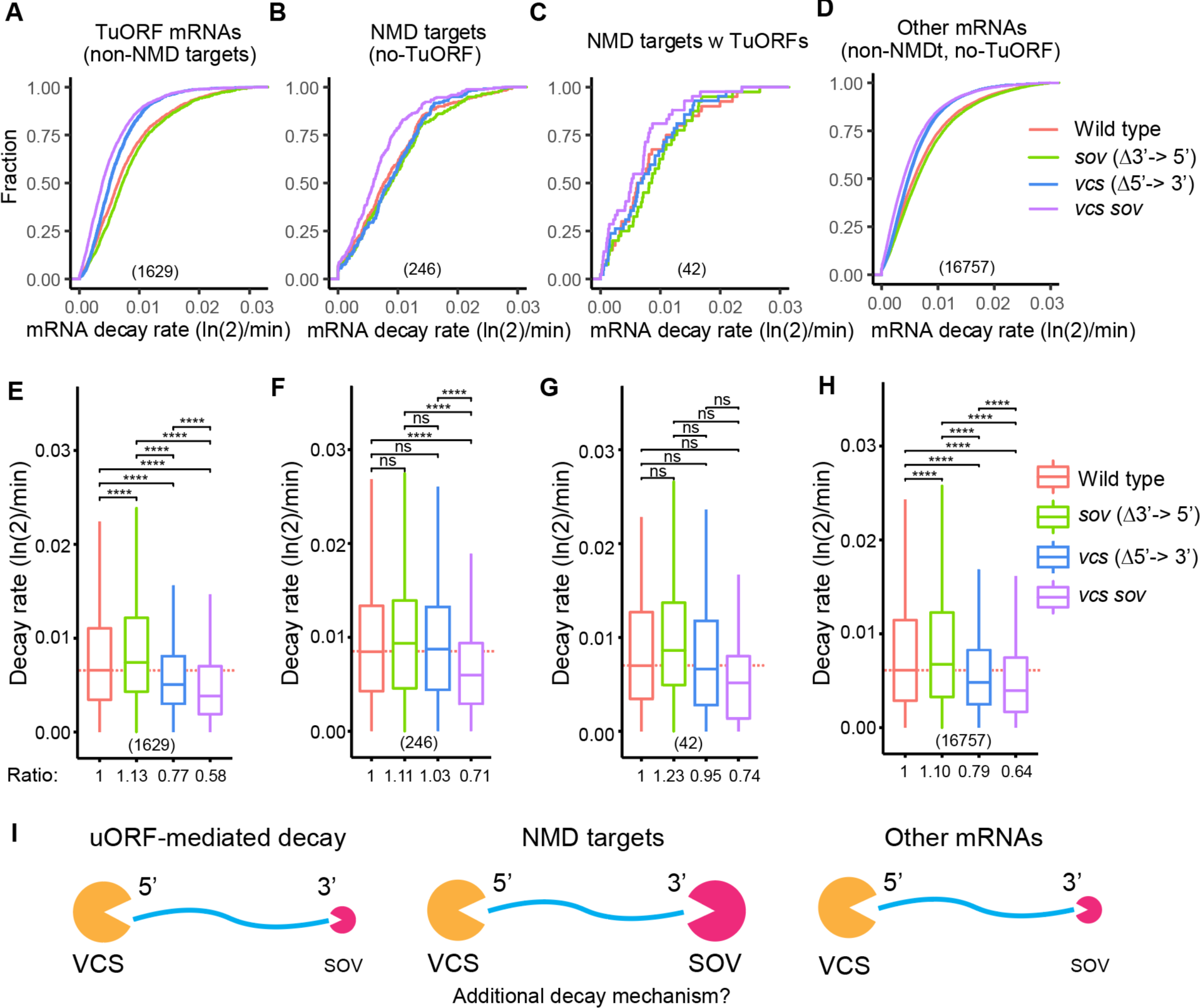
TuORF mRNAs and NMD targets are degraded through different mRNA decay mechanisms. Cumulative plots (A-D) and boxplots (E-H) showing the mRNA decay rates for NMD targets and TuORF mRNAs in wild type and mutants defective in either 5’ to 3’ decay (*vcs-7*), 3’ to 5’ decay (*sov*), or both (*vcs sov).* The mRNA decay rates were extracted from (Sorenson et al., 2018). Because Col-0 is naturally an *sov* mutant, the wild type here is Col-0 carrying the functional *SOV* allele from Landsberg *erecta* driven by its native promoter (Zhang et al., 2010; Goeres et al., 2007). The *sov* mutant here is Col-0, and *vcs* is Col-0 with the *vsc-7* allele and the L*er SOV* transgene. The *vcs sov* double mutant is Col-0 carrying the *vcs-7* mutation (Sorenson et al., 2018). Within the boxplots, the dashed lines mark the median level of wild type; the number of genes in each category is listed below the lower whisker. The ratios underneath the plots indicate the median of each group normalized to that of wild type. The statistical significance was determined as described in the Figure 2 legend. (I) Model of mRNA decay for NMD targets, TuORF mRNAs, and other mRNAs. The orange packman indicates 5’-to-3’ decay machinery, and the hot pink packman indicates 3’-to-5’ decay machinery.

In contrast, the mRNA decay rates of NMD targets without TuORFs are only significantly lower in the *vcs/sov* double mutant (**Figure 5B, 5F**), suggesting that NMD targets can be efficiently degraded by either the 5’ to 3’ or 3’ to 5’ pathways. In other words, in the absence of either *SOV* or *VCS*, the remaining pathway could completely take over the decay of NMD targets (**Figure 5B, 5F**). Although there are fewer NMD targets containing TuORFs, which resulted in small sample size and statistical insignificance (**Figure 5C, 5G**), the decay rates in this group of genes in the four genetic backgrounds show trends similar to other NMD targets without TuORFs (**Figure 5B, 5F**).

Next, we compared the decay rates of the four gene groups within each genetic background (**Figure S10)**. In all four genetic backgrounds, NMD targets with or without TuORFs consistently have higher decay rates than TuORF mRNAs and other mRNAs (**Figure S10A-H**). This pattern is still persistent in the *vcs/sov* double mutant **(Figure S10D, S10H)**, suggesting additional decay pathway(s) is involved in degrading the NMD targets in the absence of both VCS and SOV.

Taken together, our results suggest that TuORF mRNAs and NMD targets are preferentially degraded through different decay pathways. Furthermore, NMD targets consistently have higher decay rates than TuORF mRNAs and appear to be regulated by additional pathway(s) other than VCS and SOV (see the model in **Figure 5I**).

## DISCUSSION

In this study, we identified Arabidopsis TuORFs using improved Ribo-seq data and systematically examined their roles in gene expression. While our results support the expected roles of uORFs in repressing mORF translation and mRNA stability, our study reveals distinct features and regulation of uORFs in plants compared with animals.

Our observations that plant TuORF genes are associated with higher expression levels (**Figure 2**), encode larger proteins (**Figure S7B**), and are enriched for evolutionarily conserved domains (**Table S2**) raised an interesting question as to whether older genes are more likely to be controlled at the protein levels through mRNA degradation and translation via uORFs. Higher mRNA abundance may allow TuORF mRNAs to be highly translated under certain conditions that require a high level of specific proteins. For example, it has been suggested that derepression of downstream translation is a general mechanism of uORF-mediated expression under stress (Andreev et al., 2018). Thus, TuORFs could enable fast and dynamic regulation of mORF translation. Future work monitoring the translation of TuORFs and their mORFs under different conditions will help decipher the regulation and physiological roles of specific TuORFs.

If TuORFs trigger NMD in Arabidopsis, we would expect that TuORF mRNAs behave similarly to NMD targets. In this report, we presented six observations suggesting that Arabidopsis TuORF mRNAs and NMD targets are two distinct groups of transcripts: (1) NMD targets, but not TuORF mRNAs, are upregulated in various NMD mutants (**Figure 3**). (2) TuORF mRNAs have significantly higher transcript levels, while the NMD targets have lower transcript levels than other genes (**Figure 4A and 4E**). (3) NMD and TuORFs additively repress translation, suggesting they repress translation through separate mechanisms (**Figure 4B, 4F**). (4) While NMD targets are efficiently degraded through both 5’ to 3’ and 3’ to 5’ decay, TuORF mRNAs, like other genes, mainly rely on 5’ to 3’ decay (**Figure 5**). (5) The decay rates of NMD targets are consistently higher than those of TuORF mRNAs (**Figure S10)**. (6) NMD targets also appear to be degraded by additional pathway(s) in the absence of both *VCS* and *SOV* (**Figure S10H**). These differences between TuORF mRNAs and NMD targets suggest that NMD is not a major mechanism regulating TuORF mRNAs.

The fact that Arabidopsis is missing key NMD factors found in animals, including SMG1, SMG5, and SMG6, suggests significant variations in the NMD mechanisms between plants and animals. In particular, SMG1 is the kinase responsible for UPF1 phosphorylation required for substrate recognition in animal NMD (He and Jacobson, 2015; Kurosaki et al., 2019). The lack of SMG1 and the use of an alternative kinase for Arabidopsis UPF1 phosphorylation (Kerényi et al., 2013) may lead to a difference in substrate recognition: while TuORF mRNAs are recognized as NMD targets in animals, plant TuORF mRNAs are not.

Despite the difference in NMD components in plants and animals, our findings that Arabidopsis NMD targets are degraded through both 5’ to 3’ decay and 3’ to 5’ decay is consistent with the two-directional decay of NMD targets in animals. Note that plant SOV is the homolog of DIS3L2, the exonuclease responsible for 3’ to 5’ decay of NMD targets in animals (Chang et al., 2013; Lubas et al., 2013). Two-directional decay could be advantageous to quickly remove aberrant mRNAs from the cytoplasm to prevent further translation and facilitate the recycling of NMD machinery.

It is well established that there is an important interplay between translation efficiency and mRNA stability. Two mechanisms have been proposed to explain this phenomenon: the “stalled ribosome-triggered decay model,” which predicts that ribosome stalling or slowing promotes mRNA degradation, and the “translation factor-protected model,” which predicts that active translation promotes mRNA stability because the translation initiation factors protect the mRNA from the decapping complex (Presnyak et al., 2015; Radhakrishnan et al., 2016; Beelman and Parker, 1994; LaGrandeur and Parker, 1999; Schwartz and Parker, 2000; Chan et al., 2018). Consistent with the first model, several CPuORF-containing mRNAs in Arabidopsis were reported to have strong ribosome stalling patterns, and the endonucleolytic cleavages upstream of the stalled ribosomes were observed (Hou et al., 2016; Yu et al., 2016). While most of the TuORFs we identified did not display clear stalling patterns (e.g., **Figure 1F-G, S1B, S2A-N**), it remains possible that ribosomes have lower translation speeds, on average, during uORF translation. Because ribosomes will quickly approach termination after initiation due to the short length of uORFs, and because both initiation and termination are major time-consuming steps in translation (Shah et al., 2013; Lawson et al., 2021), the average translation speed per codon for uORFs is likely to be slower than that for longer ORFs. Alternatively, if cells mainly monitor the translation efficiency of the mORF, then the poorer mORF translation caused by uORFs could explain the faster degradation of TuORF mRNAs predicted by this model. In contrast, since ribosomes actively engage with TuORFs, the “translation factor-protected” model does not fit TuORF mRNAs, especially because their degradation relies on decapping and 5’ to 3’ decay in plants. Further investigation into TuORF-mediated mRNA degradation will help elucidate the role of uORFs in regulating mRNA stability.

## METHODS

### Plant growth conditions and lysate preparation

Arabidopsis Col-0 seeds were surface sterilized with 70% ethanol for 5 min, followed by 33% bleach and 0.03% Tween 20 for 10 min, then rinsed with sterile water 5 times. The seeds were imbibed at 4°C in the dark for 2 days, then grown hydroponically in sterile liquid media (2.15 g/L Murashige and Skoog salt, 1% sucrose, 0.5 g/L MES, pH 5.7) while shaking at 85 rpm under a 16-h light (75–80 μmol m^−2^·s^−1^ from cool white fluorescent bulbs) and 8-h dark cycle at 22°C for 7 days. At Zeitgeber time 4 (4 h after lights on), DMSO corresponding to 0.1% of the media volume was added to the media (these were mock samples of our large-scale experiment). After 20 and 60 min, three biological replicates (∼300 plants per sample) were harvested at each time point and immediately flash frozen in liquid nitrogen.

Plant lysates were prepared as previously described (Hsu et al., 2016). Briefly, per 0.1 g of ground tissue powder was resuspended in 400 µL of lysis buffer (100 mM Tris-HCl [pH 8], 40 mM KCl, 20 mM MgCl2, 2% [v/v] polyoxyethylene [10] tridecyl ether [Sigma, P2393], 1% [w/v] sodium deoxycholate [Sigma, D6750], 1 mM dithiothreitol, 100 µg/mL cycloheximide [Sigma, C4859], 100 µg/mL chloramphenicol [Sigma R4408], and 10 units/mL DNase I [Epicenter, D9905K]). The lysates were spun at 3,000 g for 3 min, and the supernatant was transferred to a new tube and subsequently centrifuged at 20,000 g for 10 min. The supernatant was transferred to a new tube and the RNA concentration was determined with 10x dilutions using the Qubit RNA HS assay (Thermo Fisher Scientific; Q32852). Aliquots of 100 µL and 200 µL of the lysates were made, and they were flash frozen in liquid nitrogen and stored at -80°C until further processing.

### Ribo-seq library construction

Ribosome footprints were obtained using 200 µL of the lysates described above, and sequencing libraries were constructed according to our previous method (Hsu et al., 2016) with the following modifications: briefly, after RNase I digestion (50 units nuclease per 40 µg of RNA; the nuclease was included in TruSeq Mammalian Ribo Profile Kit, Illumina, RPHMR12126), adding 15 µl of SUPERase-IN (Invitrogen, AM2696), and passing through a size exclusion column (Illustra MicroSpin S-400 HR Columns; GE Healthcare; 27-5140-01), RNA > 17 nt was isolated with the RNA Clean & Concentrator-25 kit (Zymo Research, R1017) and separated on 15% urea-TBE gels (Invitrogen, EC68852BOX). Gel slices between 28 and 30 nt were isolated, and the RNAs were purified as previously described. Next, rRNA depletion was performed using RiboZero Plant Leaf kit (Illumina, MRZPL1224) in one quarter of the recommended reaction volume. The major change we made here was to gel-purify the ribosome footprints prior to the rRNA depletion. In our previous protocol, we purified the RNA twice (first >17 nt and then <200 nt) to enrich ribosome footprints before the rRNA depletion and gel purification. We reasoned that some footprints might be lost during the two RNA purification steps. In addition, the rRNA depletion step requires a limited amount of input RNA according to the manufacturer’s recommendations. Therefore, we expected modifying the order of these steps could maximize the input and minimize the footprint loss during the processes. Ribo-seq libraries were then constructed using the TruSeq Mammalian Ribo Profile Kit (illumina, RPHMR12126) as previously described with nine cycles of PCR amplification. Libraries with equal molarity were pooled and sequenced on a Hi-Seq 4000 sequencer using single-end 50-bp sequencing.

### RNA-seq library construction

Total RNA greater than 200 nt was purified from 100 µL of the lysates described above using the RNA Clean & Concentrator-25 kit (Zymo Research, R1017) as previously described (Hsu et al., 2016). RNA integrity was evaluated using a Bioanalyzer (Agilent) RNA pico chip, and RNA integrity numbers (RINs) ranging from 7.2 to 7.7 were observed among the samples. A total of 4 µg of RNA per sample was subjected to rRNA depletion using the RiboZero Plant Leaf kit (Illumina, MRZPL1224) following the manufacturer’s recommendations. Then, 100 ng of rRNA-depleted RNA was fragmented to around 200-nt long based on the RIN reported by the Bioanalyzer, and strand-specific sequencing libraries were made using the NEBNext Ultra II Directional RNA Library Prep Kit (New England Biolabs, E7760S) with eight cycles of amplification. Libraries of equal molarity were pooled and sequenced on a Hi-Seq 4000 using paired-end 100-bp sequencing.

### Data pre-processing and analysis

Data pre-processing and analysis were performed similarly to that previously described (Hsu et al., 2016), except that the Araport11 annotation was used in this study. Briefly, for Ribo-seq libraries, the adaptor (AGATCGGAAGAGCACACGTCT) was clipped with fastx_clipper (FASTX toolkit v0.0.14) (http://hannonlab.cshl.edu/fastx_tool-kit/). For both RNA-seq and Ribo-seq, we used Bowtie2 (v2.3.4.1) (Langmead and Salzberg, 2012) to remove rRNA/tRNA/snRNA/snoRNA sequences. Both the RNA-seq and Ribo-seq reads were mapped to the transcriptome with STAR aligner (Dobin et al., 2013) (RNA-seq parameters: -- outFilterMismatchNmax 2 --outFilterMultimapNmax 20 --outFilterType BySJout --alignSJoverhangMin 8 --alignSJDBoverhangMin 2; all parameters used for Ribo-seq were the same except --outFilterMismatchNmax 1). We then used the bam files from both RNA-seq and Ribo-seq for RiboTaper (v1.3.1a) (Calviello et al., 2016) to identify translated ORFs. The distribution of Ribo-seq reads in different genome features was calculated using Ribo-seQC. The Ribo-seq metaplot was created using the create_metaplots.bash function in RiboTaper.

The Ribo-seq read lengths and offsets for RiboTaper were 24, 25, 26, 27, 28 and 8, 9, 10, 11, 12, respectively. Combining both 20-min and 60-min samples, the total reads mapped to the genome for Ribo-seq and RNA-seq were 298.01 million and 180.67 million, respectively. The discovered uORFs and mORFs were extracted from the RiboTaper output ORF_max_filt file. To calculate translation efficiency, we first used STAR to map the RNA-seq and Ribo-seq reads to the CDS of annotated coding genes. The resulting bam files were used to quantify the transcripts per million (TPM) of each gene with RSEM (v1.3.1) (Li and Dewey, 2011). Then, translation efficiency was calculated by dividing the Ribo-seq TPM by the RNA-seq TPM. The root and shoot data from Hsu et al., 2016 (Hsu et al., 2016) were reanalyzed using the Araport11 annotation. The P_sites_all files from RiboTaper were processed using the following code: cut -f 1,3,6 P_sites_all | sort | uniq -c | sed -r ’s/^(*[^]+) +/\1\t/’ > output.txt to combine the read counts at each P-site. Data visualization and statistical analysis were performed in R (v4.0.3) (R Core Team (2013), 2017). In particular, the gene levels from the Ribo-seq and RNA-seq reads were visualized with RiboPlotR (Wu and Hsu, 2021).

The mRNA half-life and mRNA decay rate data were downloaded from Szabo et al. and Sorenson et al., respectively (Szabo et al., 2020; Sorenson et al., 2018). The quantitative proteomics data for Arabidopsis shoots and roots were downloaded from Song et al. (Song et al., 2018). The expression levels (TPM) used for investigating mRNA levels of uORF genes in different tissues, developmental stages, and ecotypes in Arabidopsis and in tomato were directly downloaded from the EMBO-EBI expression atlas (E-MTAB-7978: Arabidopsis tissue atlas, E-GEOD-53197: 17 Arabidopsis thaliana accessions, E-MTAB-4812: tomato root, leaf, flower [two stages], and fruit [six stages], and E-MTAB-4813: three longitudinal sections of six stages during tomato fruit development) (Papatheodorou et al., 2020). Our previous Ribo-seq and RNA-seq data for Arabidopsis roots and shoots were described in (Hsu et al., 2016); our previous Ribo-seq and RNA-seq data for tomato roots were described in (Wu et al., 2019). The published RNA-seq data for the NMD mutants *upf1* (*lba1*) *upf3-1* and its wild-type control (GEO: GSE41432), *smg7-1 pad4* and its *pad4* control (GEO: GSE55884), *upf1-3 pad4* and its *pad4* control (GEO: GSE146340), and *upf2-10* and its wild-type control (SRA: SRP056795) were downloaded, and the mRNA levels were quantified by Kalisto (Bray et al., 2016) and the TPM values were further analyzed to compare the steady-state mRNA levels in different categories of genes.

## AVAILABILITY OF DATA AND MATERIALS

All raw and processed sequencing data generated in this study have been submitted to the NCBI Gene Expression Omnibus (GEO; https://www.ncbi.nlm.nih.gov/geo/) under accession number GSE183264. The genomic locations of TuORFs are available on TAIR genome browser (https://www.arabidopsis.org). Codes for this study are available at https://github.com/hsinyenwu/TuORF_vs_NMD_2022.

## AUTHOR CONTRIBUTIONS AND ACKNOWLEDGEMENTS

HLW and PYH designed the research, PYH performed the sequencing experiments, HLW analyzed the sequencing data, HLW and PYH interpreted the data, and HLW and PYH wrote the paper. We thank Robert Last and Melissa Lehti-Shiu for critical reading and helpful comments on the manuscript. This work used the Vincent J. Coates Genomics Sequencing Laboratory at UC Berkeley, supported by an NIH S10 OD018174 Instrumentation Grant. This work was supported by a National Science Foundation grant (2051885) and a Michigan State University startup grant to PYH.

## SUPPLEMENTARY DATA

Figure S1. Correlations among samples and comparisons between our previous and current datasets

Figure S2. Examples of TuORFs in important regulatory genes.

Figure S3. Protein abundance of the uORF-containing genes in Arabidopsis root

Figure S4. The mRNA levels and translation efficiency of TuORF mRNAs in tomato

Figure S5. mRNA levels of uORF-containing genes in different ecotypes, growth stages, and tissues in Arabidopsis and tomato

Figure S6. mRNA levels of CPuORF-containing genes compared to those of no-uORF genes

Figure S7. TuORF genes are larger than other genes

Figure S8. Lengths of the uORF peptides for NMD target and non-NMD target TuORFs

Figure S9. Lengths of UTRs and CDS for NMD targets and TuORF genes

Figure S10. The mRNA decay rates of NMD targets and TuORF-containing mRNAs in different genetic backgrounds

Table S1. Translated ORFs identified by RiboTaper

- **Table S1A-C.** Translated uORFs identified in the current dataset and our previous Arabidopsis shoot and root datasets
- **Table S1D-F.** Translated mORFs identified in the current dataset and our previous Arabidopsis shoot and root datasets

Table S2. GO-term analysis of TuORF genes

Table S3. Summary of the large-scale datasets used in this study

Table S4. NMD target gene list

- **Table S4A.** High-confidence NMD targets expressed in our data
- **Table S4B.** 49 genes identified as high-confidence NMD target TuORF genes

## REFERENCES

1. Andreev, D.E., Arnold, M., Kiniry, S.J., Loughran, G., Michel, A.M., Rachinskii, D., and Baranov, P. V. (2018). TASEP modelling provides a parsimonious explanation for the ability of a single uORF to derepress translation during the integrated stress response. eLife 7: e32563.

2. Von Arnim, A.G., Jia, Q., and Vaughn, J.N. (2014). Regulation of plant translation by upstream open reading frames. Plant Science 214: 1–12.

3. Baird, T.D., Cheng, K.C.C., Chen, Y.C., Buehler, E., Martin, S.E., Inglese, J., and Hogg, J.R. (2018). ICE1 promotes the link between splicing and nonsense-mediated mRNA decay. eLife 7.

4. Bazin, J., Baerenfaller, K., Gosai, S.J., Gregory, B.D., Crespi, M., and Bailey-Serres, J. (2017). Global analysis of ribosome-associated noncoding RNAs unveils new modes of translational regulation. Proceedings of the National Academy of Sciences of the United States of America 114: E10018–E10027.

5. Beelman, C.A. and Parker, R. (1994). Differential effects of translational inhibition in cis and in trans on the decay of the unstable yeast MFA2 mRNA. Journal of Biological Chemistry 269: 9687–9692.

6. Boudsocq, M., Barbier-Brygoo, H., and Laurière, C. (2004). Identification of nine sucrose nonfermenting 1-related protein kinases 2 activated by hyperosmotic and saline stresses in Arabidopsis thaliana. Journal of Biological Chemistry 279: 41758–41766.

7. Brar, G.A. and Weissman, J.S. (2015). Ribosome profiling reveals the what, when, where and how of protein synthesis. Nature Reviews Molecular Cell Biology 16: 651–664.

8. Bray, N.L., Pimentel, H., Melsted, P., and Pachter, L. (2016). Near-optimal probabilistic RNA-seq quantification. Nature biotechnology 34: 525–527.

9. Cai, Q., Fukushima, H., Yamamoto, M., Ishii, N., Sakamoto, T., Kurata, T., Motose, H., and Takahashi, T. (2016). The SAC51 family plays a central role in thermospermine responses in arabidopsis. Plant and Cell Physiology 57: 1583–1592.

10. Calviello, L., Mukherjee, N., Wyler, E., Zauber, H., Hirsekorn, A., Selbach, M., Landthaler, M., Obermayer, B., and Ohler, U. (2016). Detecting actively translated open reading frames in ribosome profiling data. Nature Methods 13: 165–170.

11. Calvo, S.E., Pagliarini, D.J., and Mootha, V.K. (2009). Upstream open reading frames cause widespread reduction of protein expression and are polymorphic among humans. Proceedings of the National Academy of Sciences of the United States of America 106: 7507–7512.

12. Chan, L.Y., Mugler, C.F., Heinrich, S., Vallotton, P., and Weis, K. (2018). Non-invasive measurement of mRNA decay reveals translation initiation as the major determinant of mRNA stability. eLife 7: e32536.

13. Chang, H.M., Triboulet, R., Thornton, J.E., and Gregory, R.I. (2013). A role for the Perlman syndrome exonuclease Dis3l2 in the Lin28-let-7 pathway. Nature 497: 244–248.

14. Chew, G.L., Pauli, A., and Schier, A.F. (2016). Conservation of uORF repressiveness and sequence features in mouse, human and zebrafish. Nature Communications 7: 11663.

15. Chiam, N.-C., Fujimura, T., Sano, R., Akiyoshi, N., Hiroyama, R., Watanabe, Y., Motose, H., Demura, T., and Ohtani, M. (2019). Nonsense-Mediated mRNA Decay Deficiency Affects the Auxin Response and Shoot Regeneration in Arabidopsis. Plant and Cell Physiology 60: 2000–2014.

16. Clarke, J.T., Warnock, R.C.M., and Donoghue, P.C.J. (2011). Establishing a time-scale for plant evolution. The New Phytologist 192: 266–301.

17. Cui, X., Lv, Y., Chen, M., Nikoloski, Z., Twell, D., and Zhang, D. (2015). Young genes out of the male: An insight from evolutionary age analysis of the pollen transcriptome. Molecular Plant 8: 935–945.

18. Degtiar, E., Fridman, A., Gottlieb, D., Vexler, K., Berezin, I., Farhi, R., Golani, L., and Shaul, O. (2015). The feedback control of UPF3 is crucial for RNA surveillance in plants. Nucleic Acids Research 43: 4219–4235.

19. Dever, T.E., Ivanov, I.P., and Sachs, M.S. (2020). Conserved Upstream Open Reading Frame Nascent Peptides That Control Translation. Annual Review of Genetics 54: 237–264.

20. Devlin, P.F., Patel, S.R., and Whitelam, G.C. (1998). Phytochrome E Influences Internode Elongation and Flowering Time in Arabidopsis. The Plant Cell 10: 1479–1487.

21. Deyholos, M.K., Cavaness, G.F., Hall, B., King, E., Punwani, J., Van Norman, J., and Sieburth, L.E. (2003). Varicose, a WD-domain protein, is required for leaf blade. Development 130: 6577–6588.

22. Dobin, A., Davis, C.A., Schlesinger, F., Drenkow, J., Zaleski, C., Jha, S., Batut, P., Chaisson, M., and Gingeras, T.R. (2013). STAR: ultrafast universal RNA-seq aligner. Bioinformatics 29: 15–21.

23. Drechsel, G., Kahles, A., Kesarwani, A.K., Stauffer, E., Behr, J., Drewe, P., Rätsch, G., and Wachter, A. (2013). Nonsense-mediated decay of alternative precursor mRNA splicing variants is a major determinant of the Arabidopsis steady state transcriptome. The Plant Cell 25: 3726–3742.

24. Ellis, C.M., Nagpal, P., Young, J.C., Hagen, G., Guilfoyle, T.J., and Reed, J.W. (2005). AUXIN RESPONSE FACTOR1 and AUXIN RESPONSE FACTOR2 regulate senescence and floral organ abscission in Arabidopsis thaliana. Development 132: 4563–4574.

25. Filichkin, S.A., Cumbie, J.S., Dharmawardhana, P., Jaiswal, P., Chang, J.H., Palusa, S.G., Reddy, A.S.N., Megraw, M., and Mockler, T.C. (2015). Environmental stresses modulate abundance and timing of alternatively spliced circadian transcripts in Arabidopsis. Molecular Plant 8: 207–227.

26. Gloggnitzer, J., Akimcheva, S., Srinivasan, A., Kusenda, B., Riehs, N., Stampfl, H., Bautor, J., Dekrout, B., Jonak, C., Jiménez-Gómez, J.M., Parker, J.E., and Riha, K. (2014). Nonsense-mediated mRNA decay modulates immune receptor levels to regulate plant antibacterial defense. Cell Host and Microbe 16: 376–390.

27. Goeres, D.C., Van Norman, J.M., Zhang, W., Fauver, N.A., Spencer, M. Lou, and Sieburth, L.E. (2007). Components of the Arabidopsis mRNA decapping complex are required for early seedling development. The Plant Cell 19: 1549–1564.

28. Harnett, D., Meerdink, E., Calviello, L., Sydow, D., and Ohler, U. (2021). Genome-Wide Analysis of Actively Translated Open Reading Frames Using RiboTaper/ORFquant. Methods in Molecular Biology 2252: 331–346.

29. Hayden, C.A. and Jorgensen, R.A. (2007). Identification of novel conserved peptide uORF homology groups in Arabidopsis and rice reveals ancient eukaryotic origin of select groups and preferential association with transcription factor-encoding genes. BMC Biology 5: 32.

30. He, F. and Jacobson, A. (2015). Nonsense-Mediated mRNA Decay: Degradation of Defective Transcripts Is only Part of the Story. Annual Review of Genetics 49: 339–366.

31. Hinnebusch, A.G., Ivanov, I.P., and Sonenberg, N. (2016). Translational control by 5′- untranslated regions of eukaryotic mRNAs. Science 352: 1413–1416.

32. Van Der Horst, S., Snel, B., Hanson, J., and Smeekens, S. (2019). Novel pipeline identifies new upstream ORFs and non-AUG initiating main ORFs with conserved amino acid sequences in the 5′ leader of mRNAs in Arabidopsis thaliana. RNA 25: 292–304.

33. Hou, C.-Y., Lee, W.-C., Chou, H.-C., Chen, A.-P., Chou, S.-J., and Chen, H.-M. (2016). Global analysis of truncated RNA ends reveals new insights into ribosome stalling in plants. The Plant Cell 28: 2398–2416.

34. Hsu, P.Y., Calviello, L., Wu, H.-Y.L., Li, F.-W., Rothfels, C.J., Ohler, U., and Benfey, P.N. (2016). Super-resolution ribosome profiling reveals unannotated translation events in Arabidopsis. Proceedings of the National Academy of Sciences of the United States of America 113: E7126–E7135.

35. Hu, Q., Merchante, C., Stepanova, A.N., Alonso, J.M., and Heber, S. (2016). Genome-Wide Search for Translated Upstream Open Reading Frames in Arabidopsis Thaliana. IEEE Transactions on Nanobioscience 15: 150–159.

36. Huq, E. and Quail, P.H. (2002). PIF4, a phytochrome-interacting bHLH factor, functions as a negative regulator of phytochrome B signaling in Arabidopsis. EMBO Journal 21: 2441– 2450.

37. Hurt, J.A., Robertson, A.D., and Burge, C.B. (2013). Global analyses of UPF1 binding and function reveal expanded scope of nonsense-mediated mRNA decay. Genome Research 23: 1636–1650.

38. Imai, A., Hanzawa, Y., Komura, M., Yamamoto, K.T., Komeda, Y., and Takahashi, T. (2006). The dwarf phenotype of the Arabidopsis acl5 mutant is suppressed by a mutation in an upstream ORF of a bHLH gene. Development 133: 3575–3585.

39. Ingolia, N.T., Ghaemmaghami, S., Newman, J.R.S., and Weissman, J.S. (2009). Genome- wide analysis in vivo of translation with nucleotide resolution using ribosome profiling. Science 324: 218–223.

40. Isken, O., Kim, Y.K., Hosoda, N., Mayeur, G.L., Hershey, J.W.B., and Maquat, L.E. (2008). Upf1 Phosphorylation Triggers Translational Repression during Nonsense-Mediated mRNA Decay. Cell 133: 314–327.

41. Jeong, H.J., Kim, Y.J., Kim, S.H., Kim, Y.H., Lee, I.J., Kim, Y.K., and Shin, J.S. (2011). Nonsense-mediated mRNA decay factors, UPF1 and UPF3, contribute to plant defense. Plant and Cell Physiology 52: 2147–2156.

42. Johnstone, T.G., Bazzini, A.A., and Giraldez, A.J. (2016). Upstream ORFs are prevalent translational repressors in vertebrates. The EMBO Journal 35: 706–723.

43. Jorgensen, R.A. and Dorantes-Acosta, A.E. (2012). Conserved peptide upstream open reading frames are associated with regulatory genes in angiosperms. Frontiers in Plant Science 3: 191.

44. Juntawong, P., Girke, T., Bazin, J., and Bailey-Serres, J. (2014). Translational dynamics revealed by genome-wide profiling of ribosome footprints in Arabidopsis. Proceedings of the National Academy of Sciences of the United States of America 111: E203–E212.

45. Kalyna, M. et al. (2012). Alternative splicing and nonsense-mediated decay modulate expression of important regulatory genes in Arabidopsis. Nucleic Acids Research 40: 2454–2469.

46. Kerényi, F., Wawer, I., Sikorski, P.J., Kufel, J., and Silhavy, D. (2013). Phosphorylation of the N- and C-terminal UPF1 domains plays a critical role in plant nonsense-mediated mRNA decay. Plant Journal 76: 836–848.

47. Kertész, S., Kerényi, Z., Mérai, Z., Bartos, I., Pálfy, T., Barta, E., and Silhavy, D. (2006). Both introns and long 3′-UTRs operate as cis-acting elements to trigger nonsense- mediated decay in plants. Nucleic Acids Research 34: 6147–6157.

48. Kim, B.H., Cai, X., Vaughn, J.N., and Von Arnim, A.G. (2007). On the functions of the h subunit of eukaryotic initiation factor 3 in late stages of translation initiation. Genome Biology 8: R60.

49. Kozak, M. (1999). Initiation of translation in prokaryotes and eukaryotes. Gene 234: 187–208.

50. Kurihara, Y. et al. (2009). Genome-wide suppression of aberrant mRNA-like noncoding RNAs by NMD in Arabidopsis. Proceedings of the National Academy of Sciences of the United States of America 106: 2453–2458.

51. Kurihara, Y., Makita, Y., Kawashima, M., Fujita, T., Iwasaki, S., and Matsui, M. (2018). Transcripts from downstream alternative transcription start sites evade uORF-mediated inhibition of gene expression in Arabidopsis. Proceedings of the National Academy of Sciences of the United States of America 115: 7831–7836.

52. Kurihara, Y., Makita, Y., Shimohira, H., Fujita, T., Iwasaki, S., and Matsui, M. (2020). Translational landscape of protein-coding and non-protein-coding RNAs upon light exposure in Arabidopsis. Plant and Cell Physiology 61: 536–545.

53. Kurosaki, T., Li, W., Hoque, M., Popp, M.W.L., Ermolenko, D.N., Tian, B., and Maquat, L.E. (2014). A Post-Translational regulatory switch on UPF1 controls targeted mRNA degradation. Genes and Development 28: 1900–1916.

54. Kurosaki, T., Popp, M.W., and Maquat, L.E. (2019). Quality and quantity control of gene expression by nonsense-mediated mRNA decay. Nature Reviews Molecular Cell Biology 2019 20:7 20: 406–420.

55. LaGrandeur, T. and Parker, R. (1999). The cis acting sequences responsible for the differential decay of the unstable MFA2 and stable PGK1 transcipts in yeast include the context of the translational start codon. RNA 5: 420–433.

56. Langmead, B. and Salzberg, S.L. (2012). Fast gapped-read alignment with Bowtie 2. Nature Methods 9: 357–359.

57. Di Laurenzio, L., Wysocka-Diller, J., Malamy, J.E., Pysh, L., Helariutta, Y., Freshour, G., Hahn, M.G., Feldmann, K.A., and Benfey, P.N. (1996). The SCARECROW gene regulates an asymmetric cell division that is essential for generating the radial organization of the Arabidopsis root. Cell 86: 423–433.

58. Lawson, M.R., Lessen, L.N., Wang, J., Prabhakar, A., Corsepius, N.C., Green, R., and Puglisi, J.D. (2021). Mechanisms that ensure speed and fidelity in eukaryotic translation termination. Science 373: 876–882.

59. Lee, D.S.M., Park, J., Kromer, A., Baras, A., Rader, D.J., Ritchie, M.D., Ghanem, L.R., and Barash, Y. (2021). Disrupting upstream translation in mRNAs is associated with human disease. Nature Communications 2021 12:1 12: 1–14.

60. Leivar, P., Monte, E., Al-Sady, B., Carle, C., Storer, A., Alonso, J.M., Ecker, J.R., and Quail, P.H. (2008). The Arabidopsis phytochrome-interacting factor PIF7, together with PIF3 and PIF4, regulates responses to prolonged red light by modulating phyB levels. The Plant Cell 20: 337–352.

61. Li, B. and Dewey, C.N. (2011). RSEM: accurate transcript quantification from RNA-Seq data with or without a reference genome. BMC Bioinformatics 12: 323.

62. Li, Y.R. and Liu, M.J. (2020). Prevalence of alternative AUG and non-AUG translation initiators and their regulatory effects across plants. Genome Research 30: 1418–1433.

63. Lin, Y., May, G.E., Kready, H., Nazzaro, L., Mao, M., Spealman, P., Creeger, Y., and Joel McManus, C. (2019). Impacts of uORF codon identity and position on translation regulation. Nucleic Acids Research 47: 9358–9367.

64. Liu, M.J., Wu, S.H., Wu, J.F., Lin, W.D., Wu, Y.C., Tsai, T.Y., Tsai, H.L., and Wu, S.H. (2013). Translational landscape of photomorphogenic Arabidopsis. Plant Cell 25: 3699–3710.

65. Lloyd, J.P.B. and Davies, B. (2013). SMG1 is an ancient nonsense-mediated mRNA decay effector. The Plant journal : for cell and molecular biology 76: 800–810.

66. Lubas, M., Damgaard, C.K., Tomecki, R., Cysewski, D., Jensen, T.H., and Dziembowski, A. (2013). Exonuclease hDIS3L2 specifies an exosome-independent 3’-5’ degradation pathway of human cytoplasmic mRNA. The EMBO journal 32: 1855–1868.

67. Merchante, C., Brumos, J., Yun, J., Hu, Q., Spencer, K.R., Enríquez, P., Binder, B.M., Heber, S., Stepanova, A.N., and Alonso, J.M. (2015). Gene-specific translation regulation mediated by the hormone-signaling molecule EIN2. Cell 163: 684–697.

68. Mizoguchi, T., Hayashida, N., Yamaguchi-Shinozaki, K., Kamada, H., and Shinozaki, K. (1995). Two genes that encode ribosomal-protein S6 kinase homologs are induced by cold or salinity stress in Arabidopsis thaliana. FEBS Letters 358: 199–204.

69. Nagarajan, V.K., Kukulich, P.M., Von Hagel, B., and Green, P.J. (2019). RNA degradomes reveal substrates and importance for dark and nitrogen stress responses of Arabidopsis XRN4. Nucleic Acids Research 47: 9216–9230.

70. Nagpal, P., Ellis, C.M., Weber, H., Ploense, S.E., Barkawi, L.S., Guilfoyle, T.J., Hagen, G., Alonso, J.M., Cohen, J.D., Farmer, E.E., Ecker, J.R., and Reed, J.W. (2005). Auxin response factors ARF6 and ARF8 promote jasmonic acid production and flower maturation. Development 132: 4107–4118.

71. Ni, M., Tepperman, J.M., and Quail, P.H. (1998). PIF3, a phytochrome-interacting factor necessary for normal photoinduced signal transduction, is a novel basic helix-loop-helix protein. Cell 95: 657–667.

72. Nyikó, T., Sonkoly, B., Mérai, Z., Benkovics, A.H., and Silhavy, D. (2009). Plant upstream ORFs can trigger nonsense-mediated mRNA decay in a size-dependent manner. Plant Molecular Biology 71: 367–378.

73. Ohtani, M. and Wachter, A. (2019). NMD-Based Gene Regulation - A Strategy for Fitness Enhancement in Plants? Plant and Cell Physiology 60: 1953–1960.

74. Papatheodorou, I. et al. (2020). Expression Atlas update: From tissues to single cells. Nucleic Acids Research 48: D77–D83.

75. Park, Y.S., Hong, S.W., Oh, S.A., Kwak, J.M., Lee, H.H., and Nam, H.G. (1993). Two putative protein kinases from Arabidopsis thaliana contain highly acidic domains. Plant Molecular Biology 22: 615–624.

76. Presnyak, V., Alhusaini, N., Chen, Y.H., Martin, S., Morris, N., Kline, N., Olson, S., Weinberg, D., Baker, K.E., Graveley, B.R., and Coller, J. (2015). Codon optimality is a major determinant of mRNA stability. Cell 160: 1111–1124.

77. R Core Team (2013) (2017). R: A language and environment for statistical computing. R Foundation for Statistical Computing, Vienna, Austria.

78. Radhakrishnan, A., Chen, Y.H., Martin, S., Alhusaini, N., Green, R., and Coller, J. (2016). The DEAD-Box Protein Dhh1p Couples mRNA Decay and Translation by Monitoring Codon Optimality. Cell 167: 122–132.e9.

79. Raxwal, V.K. and Riha, K. (2016). Nonsense mediated RNA decay and evolutionary capacitance. Biochimica et Biophysica Acta 1859: 1538–1543.

80. Raxwal, V.K., Simpson, C.G., Gloggnitzer, J., Entinze, J.C., Guo, W., Zhang, R., Brown, J.W.S., and Riha, K. (2020). Nonsense-Mediated RNA Decay Factor UPF1 Is Critical for Posttranscriptional and Translational Gene Regulation in Arabidopsis. The Plant Cell 32: 2725–2741.

81. Rayson, S., Arciga-Reyes, L., Wootton, L., de Torres Zabala, M., Truman, W., Graham, N., Grant, M., and Davies, B. (2012). A role for nonsense-mediated mrna decay in plants: Pathogen responses are induced in arabidopsis thaliana nmd mutants. PLoS ONE 7: e31917.

82. Riehs-Kearnan, N., Gloggnitzer, J., Dekrout, B., Jonak, C., and Riha, K. (2012). Aberrant growth and lethality of Arabidopsis deficient in nonsense-mediated RNA decay factors is caused by autoimmune-like response. Nucleic Acids Research 40: 5615–5624.

83. Saul, H. et al. (2009). The upstream open reading frame of the Arabidopsis AtMHX gene has a strong impact on transcript accumulation through the nonsense-mediated mRNA decay pathway. The Plant Journal 60: 1031–1042.

84. Schwartz, D.C. and Parker, R. (2000). mRNA Decapping in Yeast Requires Dissociation of the Cap Binding Protein, Eukaryotic Translation Initiation Factor 4E. Molecular and Cellular Biology 20: 7933–7942.

85. Shah, P., Ding, Y., Niemczyk, M., Kudla, G., and Plotkin, J.B. (2013). Rate-Limiting Steps in Yeast Protein Translation. Cell 153: 1589–1601.

86. Sharrock, R.A. and Quail, P.H. (1989). Novel phytochrome sequences in Arabidopsis thaliana: structure, evolution, and differential expression of a plant regulatory photoreceptor family. Genes & Development 3: 1745–1757.

87. Sieburth, L.E. and Vincent, J.N. (2018). Beyond transcription factors: Roles of mrna decay in regulating gene expression in plants [version 1; referees: 3 approved]. F1000Research 7: 1940.

88. Silverstone, A.L., Ciampaglio, C.N., and Sun, T.P. (1998). The Arabidopsis RGA gene encodes a transcriptional regulator repressing the gibberellin signal transduction pathway. The Plant Cell 10: 155–169.

89. Song, G., Hsu, P.Y., and Walley, J.W. (2018). Assessment and Refinement of Sample Preparation Methods for Deep and Quantitative Plant Proteome Profiling. Proteomics 18: e1800220.

90. Sorenson, R.S., Deshotel, M.J., Johnson, K., Adler, F.R., and Sieburth, L.E. (2018). Arabidopsis mRNA decay landscape arises from specialized RNA decay substrates, decapping-mediated feedback, and redundancy. Proceedings of the National Academy of Sciences of the United States of America 115: E1485–E1494.

91. Strayer, C., Oyama, T., Schultz, T.F., Raman, R., Somers, D.E., Mas, P., Panda, S., Kreps, J.A., and Kay, S.A. (2000). Cloning of the Arabidopsis clock gene TOC1, an autoregulatory response regulator homolog. Science 289: 768–771.

92. Szabo, E.X., Reichert, P., Lehniger, M.K., Ohmer, M., de Francisco Amorim, M., Gowik, U., Schmitz-Linneweber, C., and Laubinger, S. (2020). Metabolic labeling of RNAs uncovers hidden features and dynamics of the Arabidopsis transcriptome. The Plant Cell 32: 871– 887.

93. Takahashi, H., Takahashi, A., Naito, S., and Onouchi, H. (2012). BAIUCAS: A novel BLAST- based algorithm for the identification of upstream open reading frames with conserved amino acid sequences and its application to the Arabidopsis thaliana genome. Bioinformatics 28: 2231–2241.

94. Tanaka, M., Sotta, N., Yamazumi, Y., Yamashita, Y., Miwa, K., Murota, K., Chiba, Y., Hirai, M.Y., Akiyama, T., Onouchi, H., Naito, S., and Fujiwara, T. (2016). The minimum open reading frame, AUG-Stop, induces boron-dependent ribosome stalling and mRNA degradation. The Plant Cell 28: 2830–2849.

95. Uchiyama-Kadokura, N., Murakami, K., Takemoto, M., Koyanagi, N., Murota, K., Naito, S., and Onouchi, H. (2014). Polyamine-responsive ribosomal arrest at the stop codon of an upstream open reading frame of the AdoMetDC1 gene triggers nonsense-mediated mRNA decay in Arabidopsis thaliana. Plant and Cell Physiology 55: 1556–1567.

96. Vaughn, J.N., Ellingson, S.R., Mignone, F., and Arnim, A. von (2012). Known and novel post-transcriptional regulatory sequences are conserved across plant families. RNA 18: 368–84.

97. Wolfa, Y.I., Novichkovb, P.S., Kareva, G.P., Koonina, E. V., and Lipmana, D.J. (2009). The universal distribution of evolutionary rates of genes and distinct characteristics of eukaryotic genes of different apparent ages. Proceedings of the National Academy of Sciences of the United States of America 106: 7273–7280.

98. Wu, H.-Y.L. and Hsu, P.Y. (2021). RiboPlotR: a visualization tool for periodic Ribo-seq reads. Plant Methods 17: 1–7.

99. Wu, H.Y.L., Song, G., Walley, J.W., and Hsu, P.Y. (2019). The tomato translational landscape revealed by transcriptome assembly and ribosome profiling. Plant Physiology 181: 367– 380.

100. Xu, G., Yuan, M., Ai, C., Liu, L., Zhuang, E., Karapetyan, S., Wang, S., and Dong, X. (2017). uORF-mediated translation allows engineered plant disease resistance without fitness costs. Nature 545: 491–494.

101. Xu, J., Yang, J.Y., Niu, Q.W., and Chua, N.H. (2006). Arabidopsis DCP2, DCP1, and VARICOSE form a decapping complex required for postembryonic development. The Plant Cell 18: 3386–3398.

102. Xu, Z., Hu, L., Shi, B., Geng, S., Xu, L., Wang, D., and Lu, Z.J. (2018). Ribosome elongating footprints denoised by wavelet transform comprehensively characterize dynamic cellular translation events. Nucleic Acids Research 46: e109.

103. Yoine, M., Ohto, M.A., Onai, K., Mita, S., and Nakamura, K. (2006). The lba1 mutation of UPF1 RNA helicase involved in nonsense-mediated mRNA decay causes pleiotropic phenotypic changes and altered sugar signalling in Arabidopsis. The Plant Journal 47: 49– 62.

104. Yu, X., Willmann, M.R., Anderson, S.J., and Gregory, B.D. (2016). Genome-wide mapping of uncapped and cleaved transcripts reveals a role for the nuclear mrna cap-binding complex in cotranslational rna decay in Arabidopsis. The Plant Cell 28: 2385–2397.

105. Zhang, H., Si, X., Ji, X., Fan, R., Liu, J., Chen, K., Wang, D., and Gao, C. (2018). Genome editing of upstream open reading frames enables translational control in plants. Nature Biotechnology 36: 894–898.

106. Zhang, H., Wang, Y., Wu, X., Tang, X., Wu, C., and Lu, J. (2021). Determinants of genome- wide distribution and evolution of uORFs in eukaryotes. Nature Communications 12: 1–17.

107. Zhang, S.H., Lawton, M.A., Hunter, T., and Lamb, C.J. (1994). atpk1, a novel ribosomal protein kinase gene from Arabidopsis. I. Isolation, characterization, and expression. Journal of Biological Chemistry 269: 17586–17592.

108. Zhang, W., Murphy, C., and Sieburth, L.E. (2010). Conserved RNaseII domain protein functions in cytoplasmic mRNA decay and suppresses Arabidopsis decapping mutant phenotypes. Proceedings of the National Academy of Sciences of the United States of America 107: 15981–15985.

109. Zinshteyn, B., Sinha, N.K., Enam, S.U., Koleske, B., and Green, R. (2021). Translational repression of NMD targets by GIGYF2 and EIF4E2. PLoS Genetics 17: e1009813.

