## Supplemental figures for "Arabidopsis uORF-containing mRNAs behave differently from NMD targets"

### SUPPLEMENTARY FIGURES

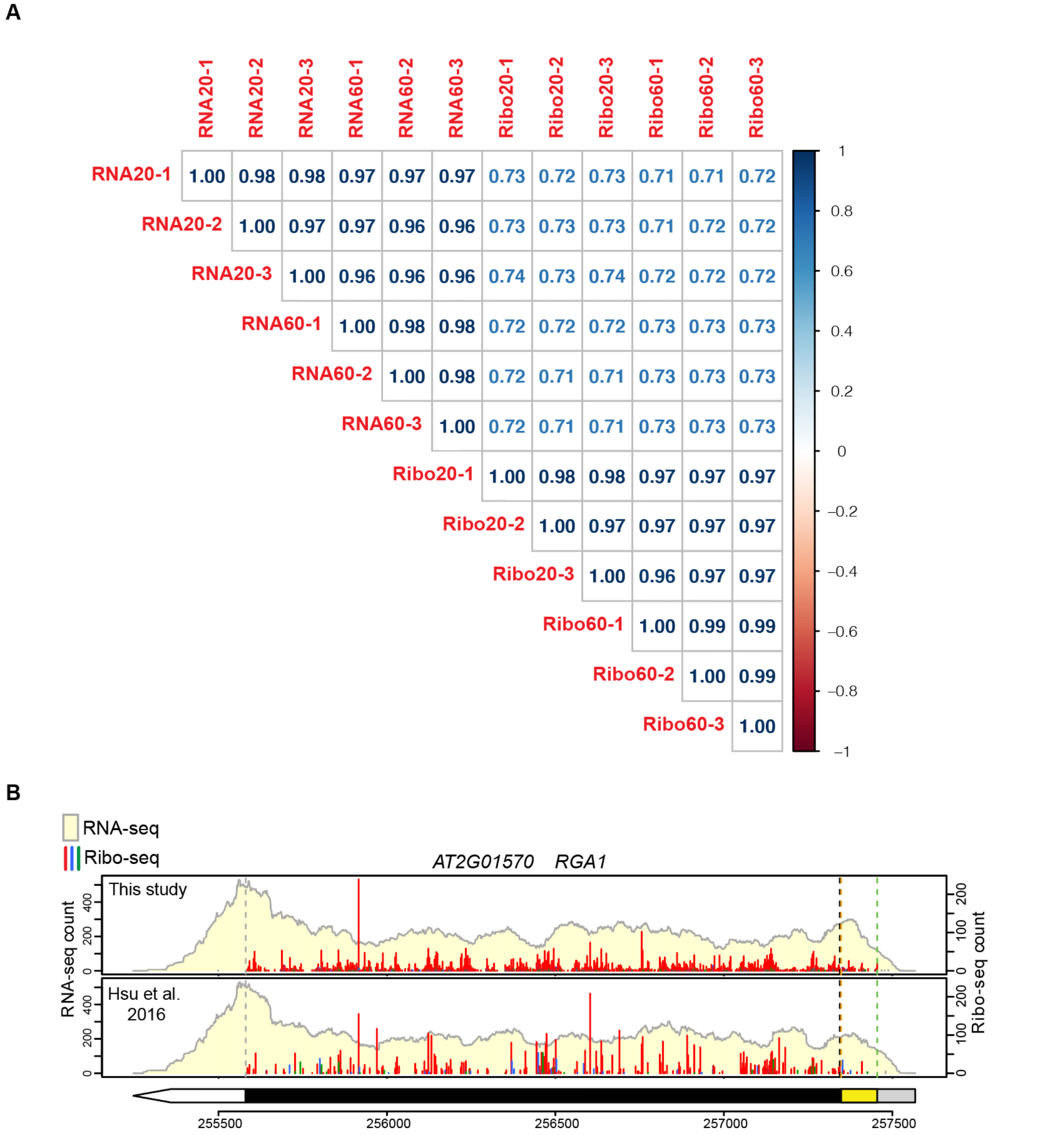


**Figure S1. Correlations among samples and comparisons between previous and current RNA-seq and Ribo-seq datasets.** (A) Pearson correlations among three biological replicates of our current RNA-seq and Ribo-seq analyses of samples harvested after 20 or 60 min of DMSO treatment (these were control samples of our large-scale experiment). TPM values in the CDSs of annotated protein-coding genes were used. (B) Higher Ribo-seq read coverage was obtained in both the uORF and mORF regions in our current dataset; *RGA1* is shown as an example. Note that the mRNA levels are similar between the previous data ((Hsu et al., 2016), shoot) and the current data. Data representations are the same as those described in the Figure 1F legend.

**Figure S2. TuORFs in known important regulatory genes**

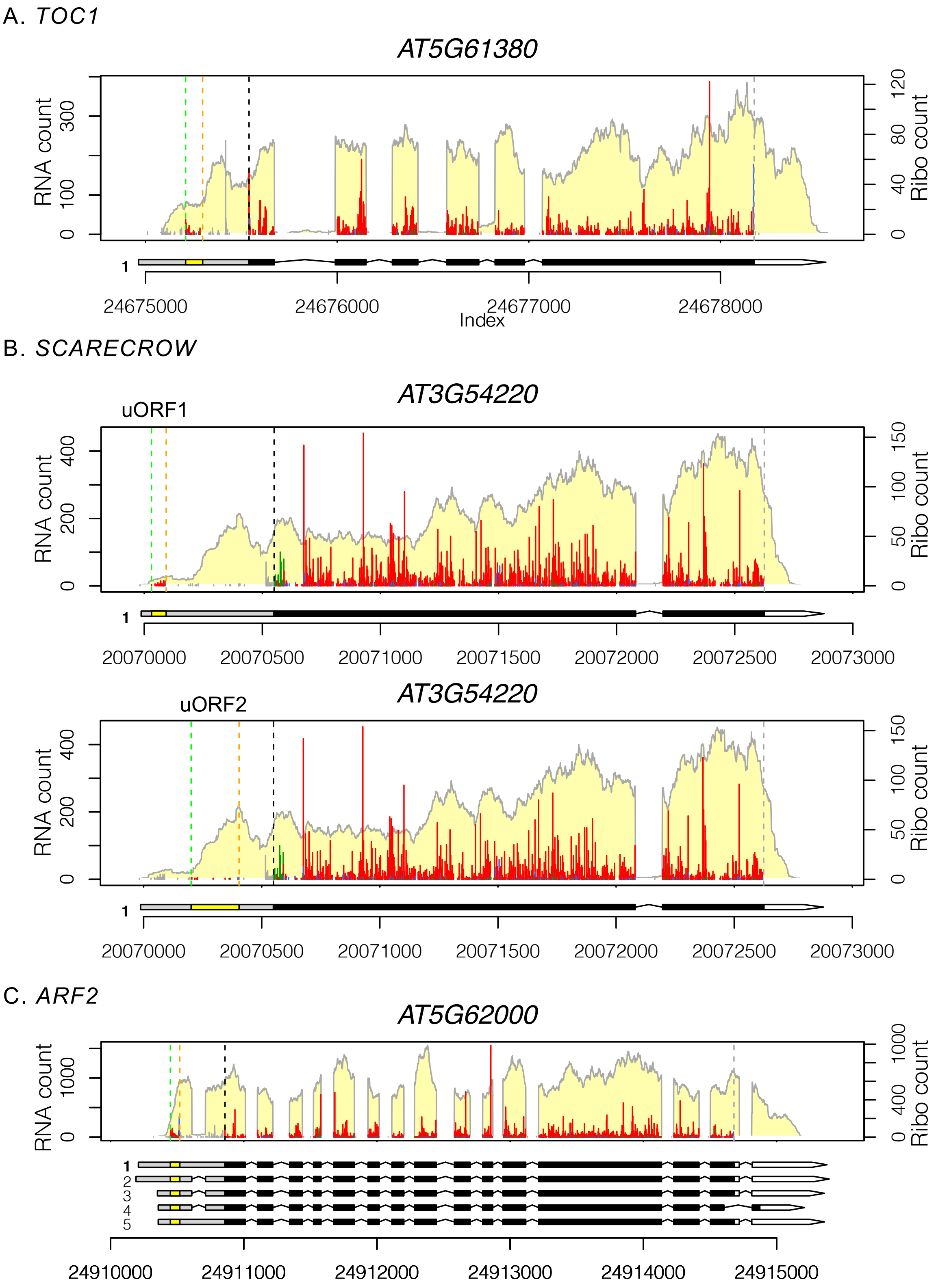

D. *ARF3*

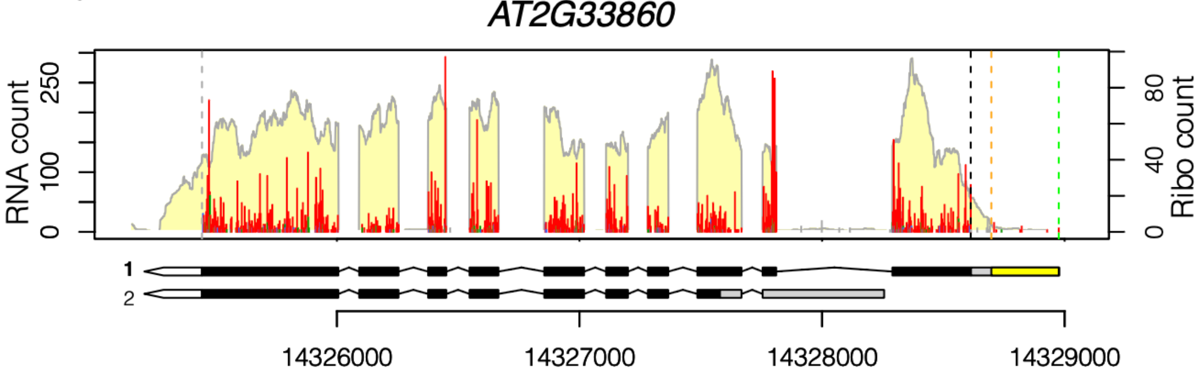
AT2G33860
RNA count
Ribo count
0
100
250
0
40
80
1
2
14326000
14327000
14328000
14329000

E. *ARF8*

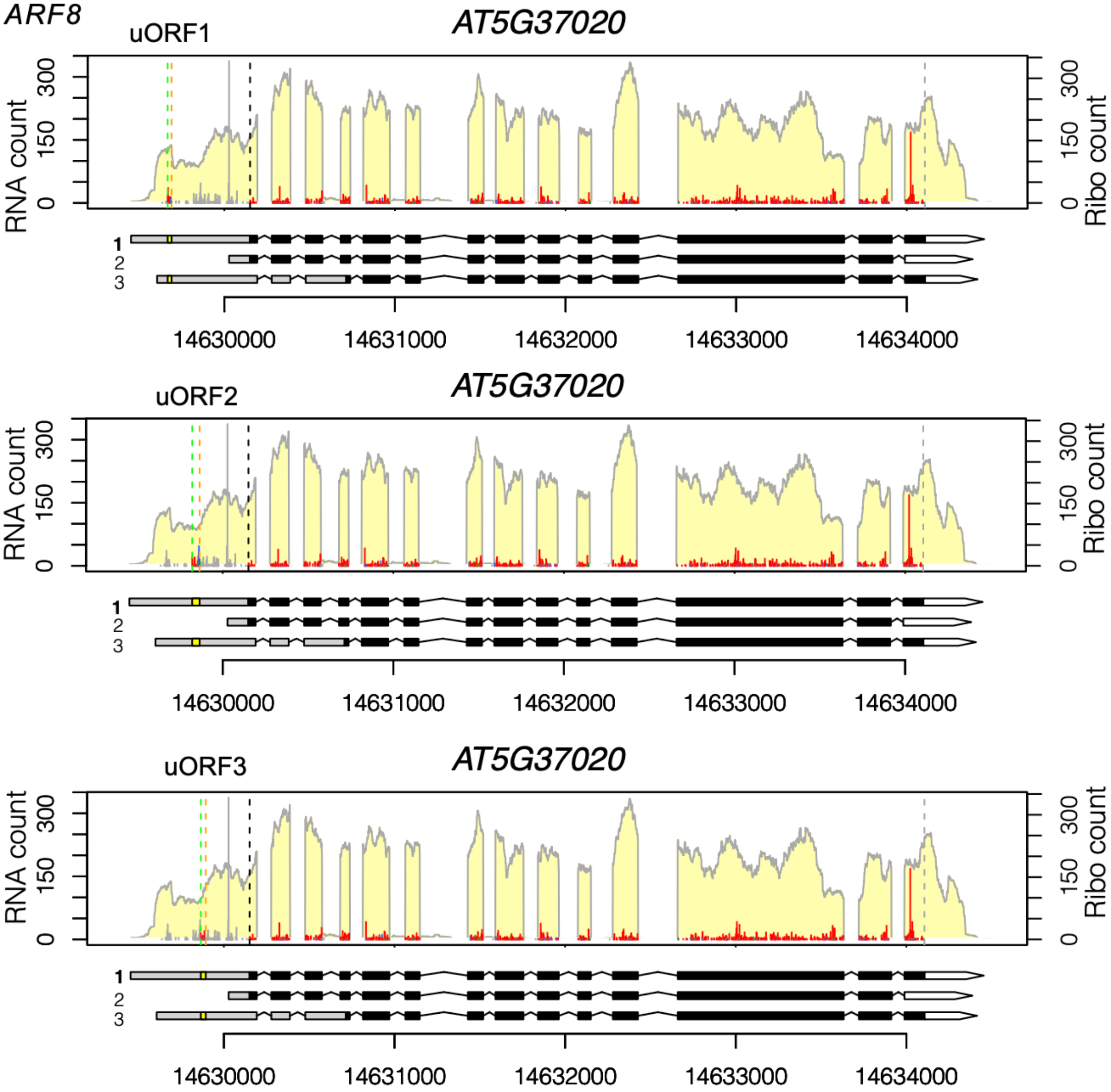
uORF1
AT5G37020
uORF2
AT5G37020
uORF3
AT5G37020
RNA count
Ribo count
0
150
300
1
2
3
14630000
14631000
14632000
14633000
14634000

E. *ARF8* (continued)

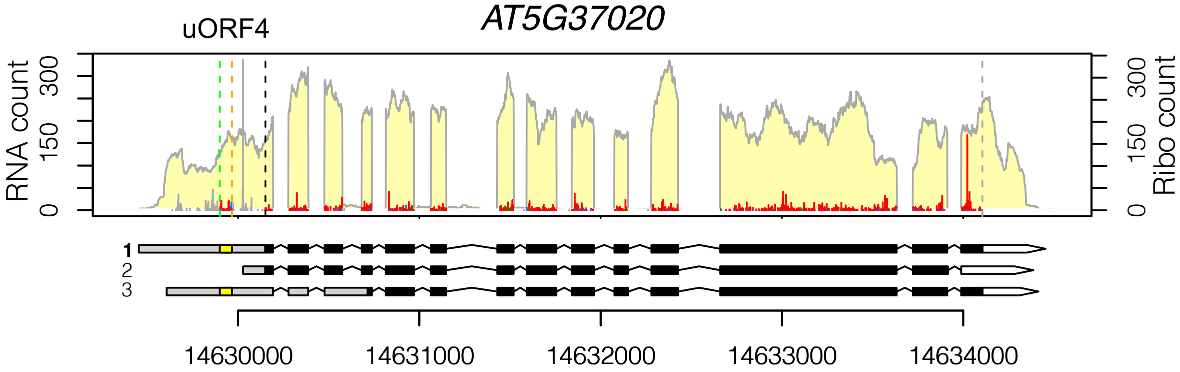


F. *Phytochrome B*

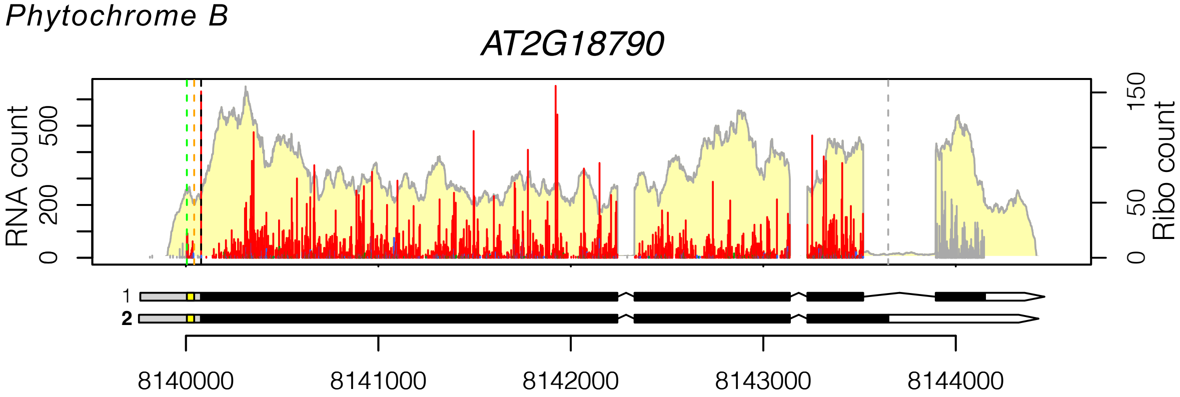


G. *Phytochrome E*

*AT4G18130*

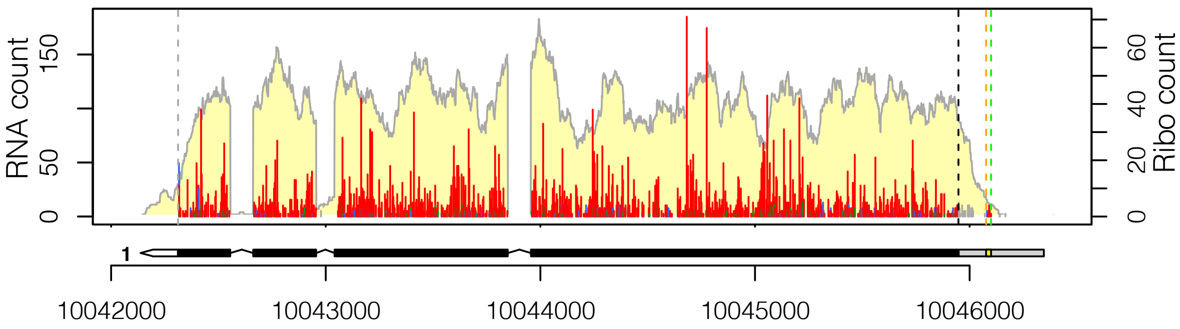


H. *PIF3*

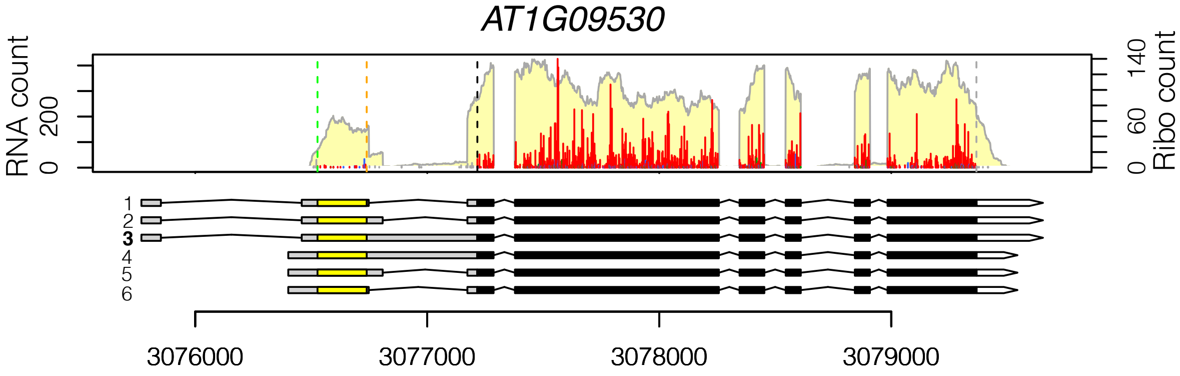

I. *PIF4*

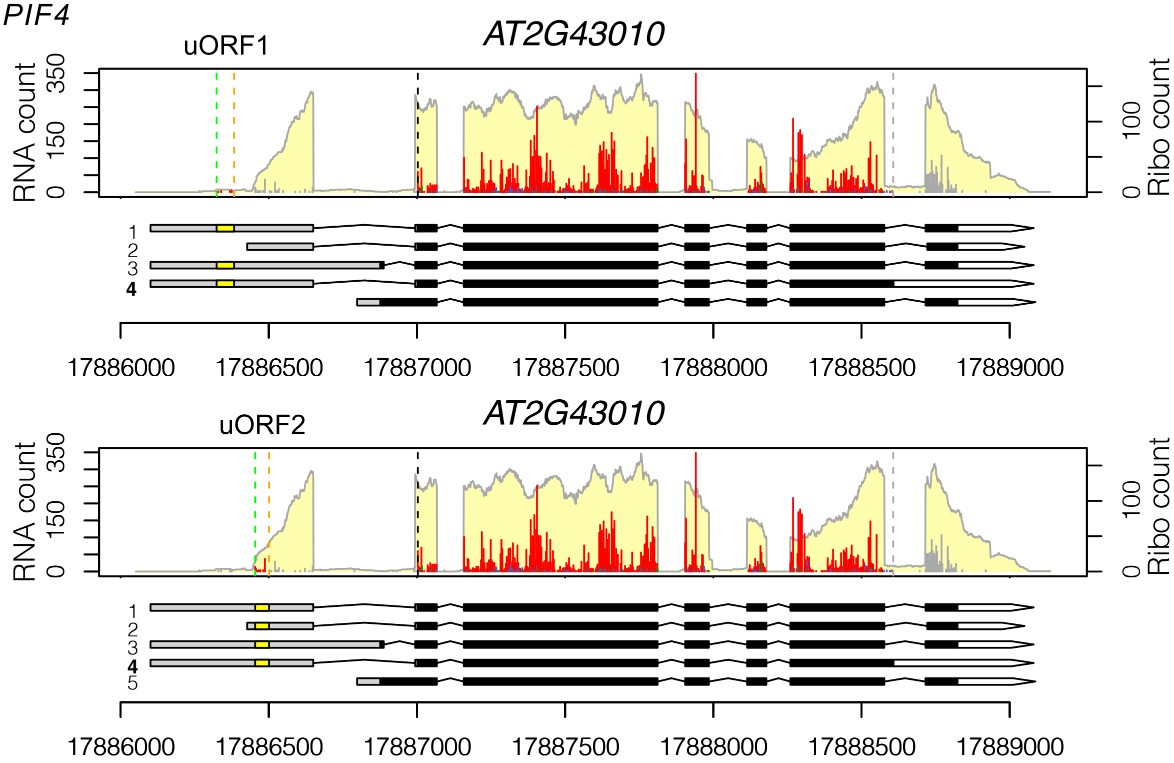


J. *PIF7*

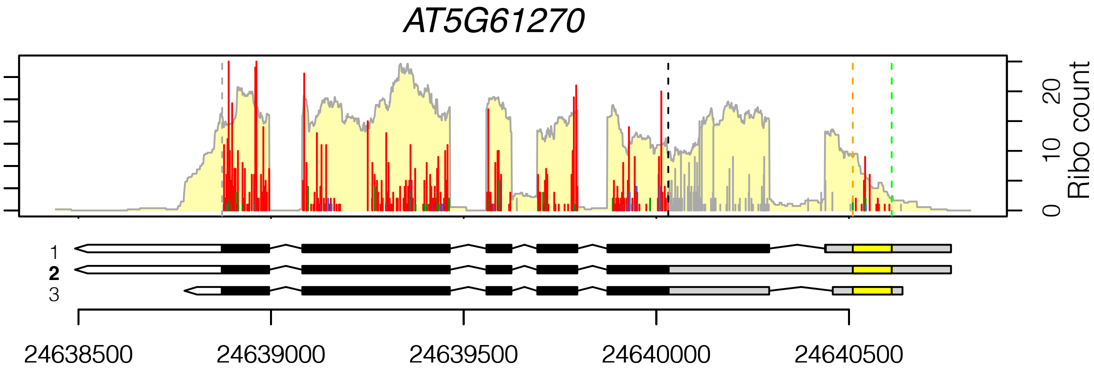


K. *S6K1*

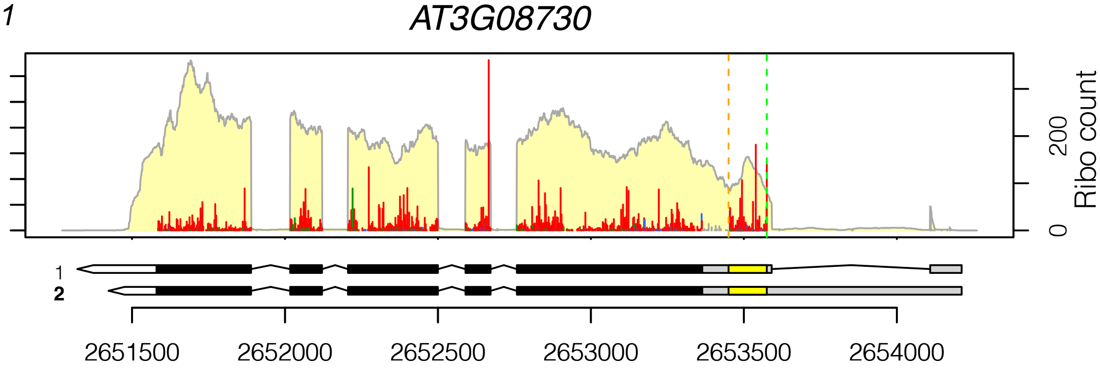

L. *S6K2*

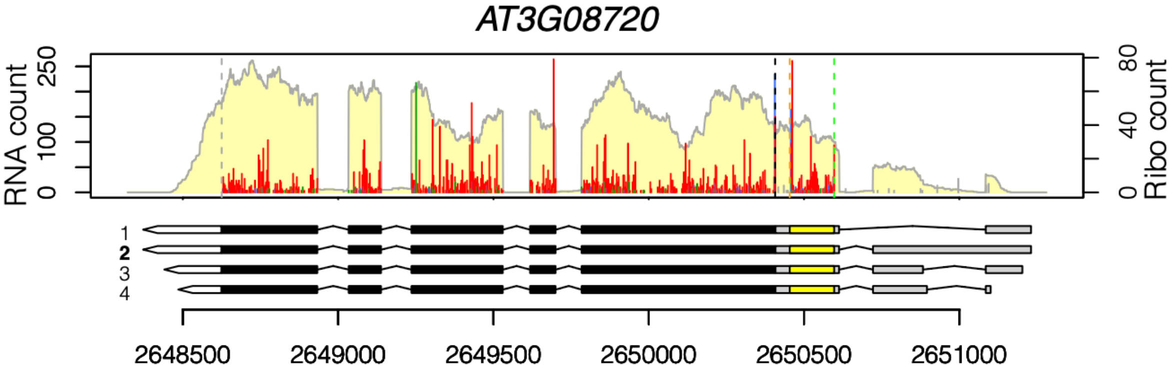


M. *SNRK2.1*

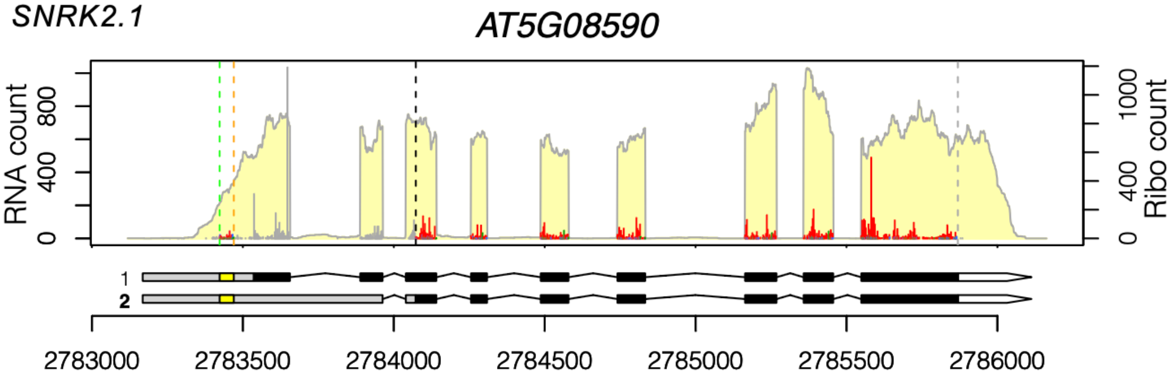


N. *SNRK2.5*

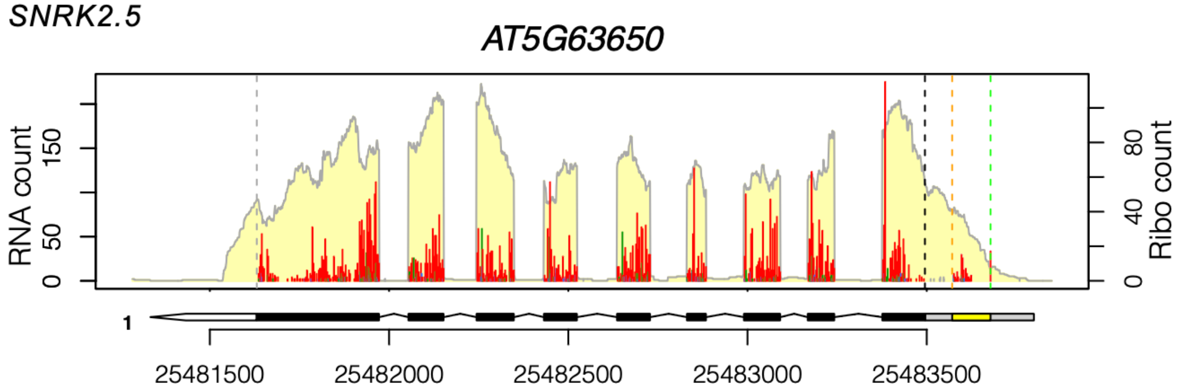


**Figure S2. Examples of TuORFs in important regulatory genes.** Data are presented as described in the Figure 1F legend. Isoform numbers are shown to the left of the gene models, and the isoform being considered is bolded. For genes that have multiple TuORFs identified (B, E, and I), the 3-nt periodicity of each TuORF is presented in a separate panel.

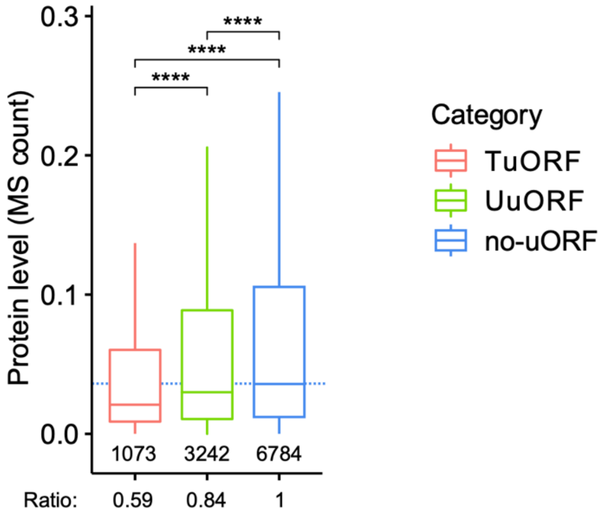


**Figure S3. Protein abundance of the uORF-containing genes in Arabidopsis roots.** The protein quantification data from Arabidopsis seedling roots were extracted from (Song et al., 2018). Within the boxplot, the dashed line marks the median level of no-uORF genes; the number of genes in each category is listed below the lower whisker. The ratios underneath the plots indicate the median of each group normalized to that of no-uORF genes. The statistical significance was determined as described in the Figure 2 legend.

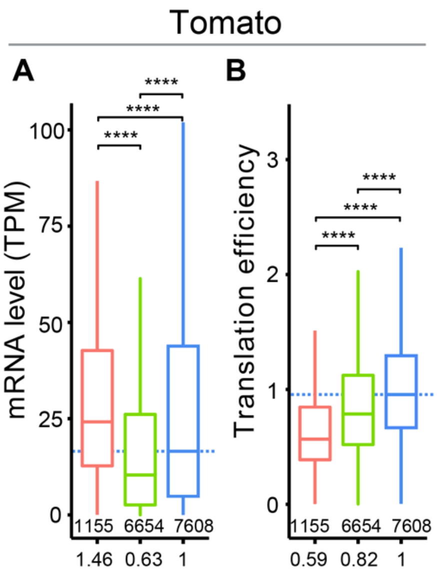


**Figure S4. Tomato TuORF mRNAs are also associated with higher mRNA abundance and lower translation efficiency.** The mRNA levels and translation efficiency of TuORF, UuORF, and no-uORF genes from a reanalysis of our previous tomato root data (Wu et al., 2019) were presented. The dashed lines mark the median level of the no-uORF genes.

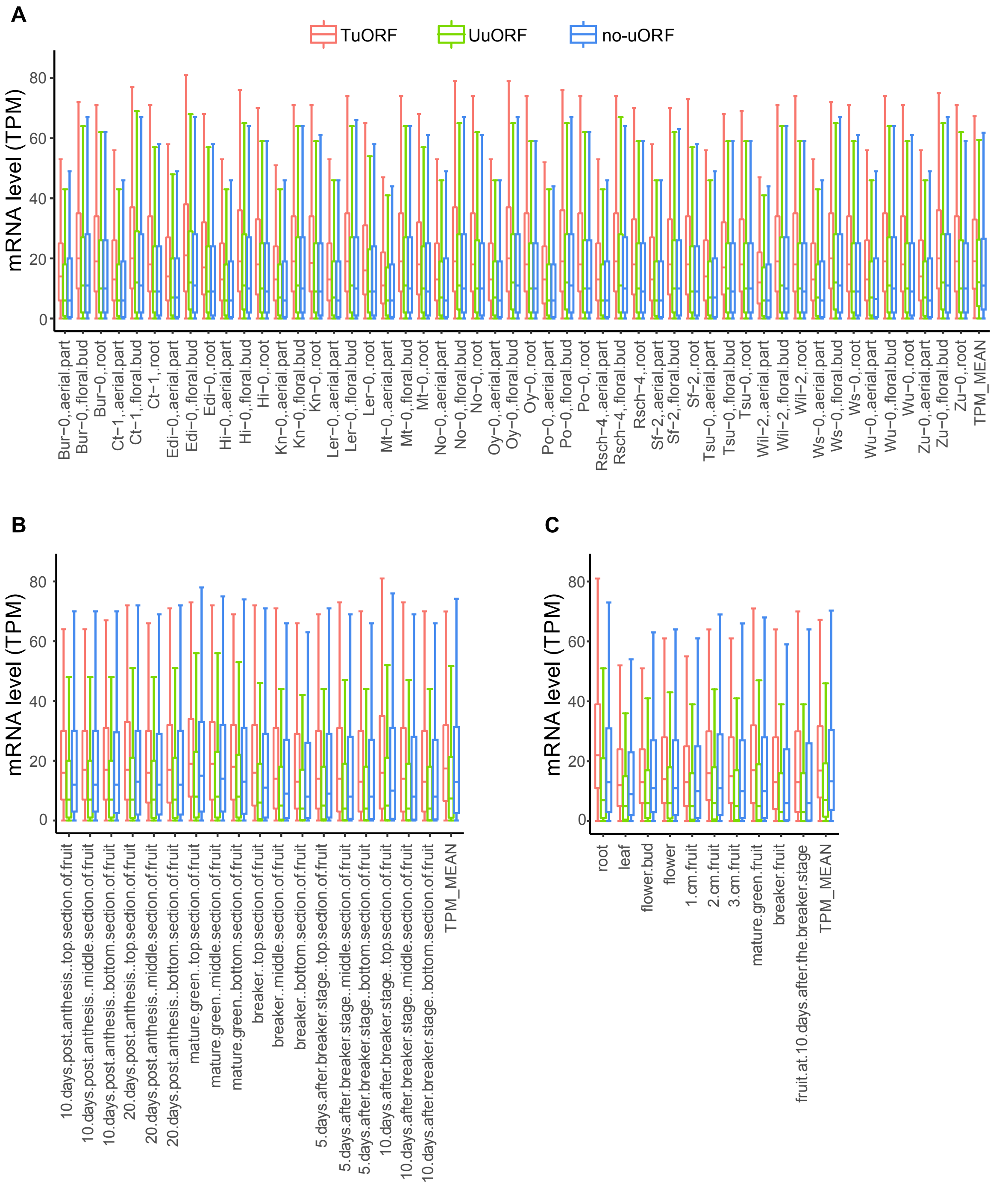


**Figure S5. mRNA levels of uORF-containing genes in different tissues, developmental stages and ecotypes in Arabidopsis and tomato.**

TuORF, UuORF, and no-uORF genes were compared.

(A) Arabidopsis roots and aerial and floral parts from different ecotypes.

(B) Various developmental stages of tomato fruit (Heinz 1706 cultivar).

(C) Roots, shoots, flowers, and fruits of tomato (Heinz 1706 cultivar).

The TPM values from RNA-seq quantification were extracted from the EMBO-EBI expression atlas (Papatheodorou et al., 2020) (see METHODS).

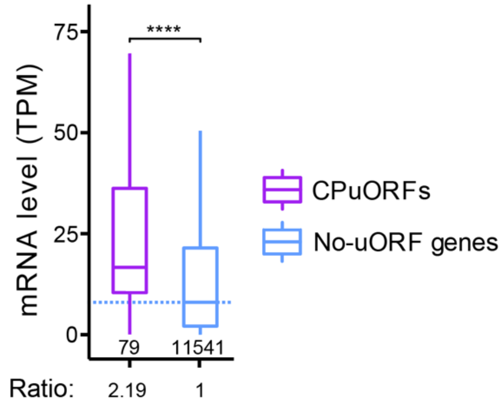


**Figure S6. mRNA levels of CPuORF-containing genes compared with no-uORF genes.** Within the boxplot, the dashed line marks the median level of no-uORF genes; the number of genes in each category is listed below the lower whisker. The ratios underneath the plots indicate the median of each group normalized to that of no-uORF genes. The statistical significance was determined as described in the Figure 2 legend.

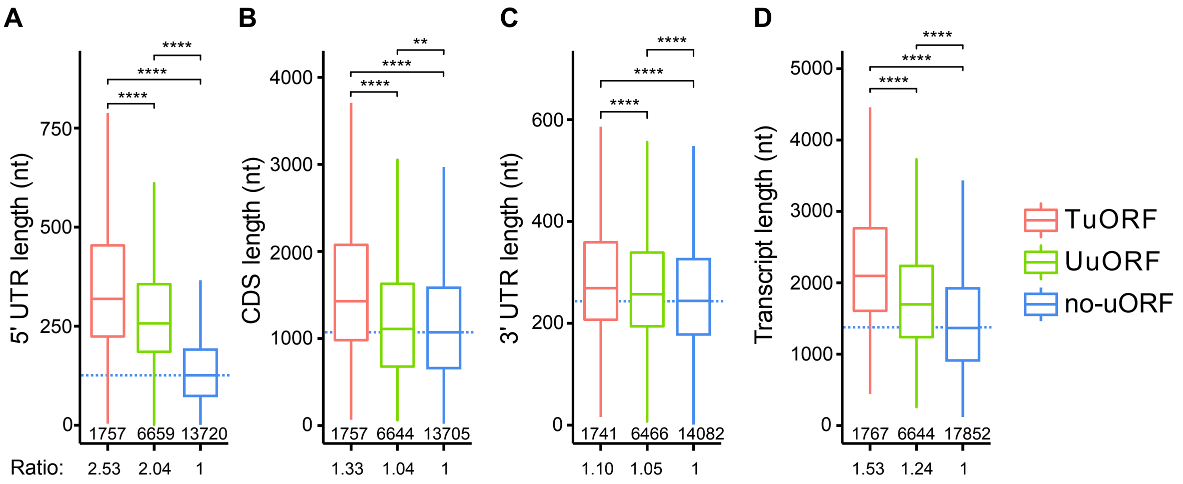


**Figure S7. TuORF genes are larger than other genes**

Lengths of the 5' UTRs (A), CDSs (B), 3' UTRs (C), and entire transcripts (D) of TuORF, UuORF, and no-uORF genes. Within the boxplots, the dashed lines mark the median level of the no-uORF genes; the number of genes in each category is listed below the lower whisker. The ratios underneath the plots indicate the median of each group normalized to that of the no-uORF genes. The statistical significance was determined as described in the Figure 2 legend.

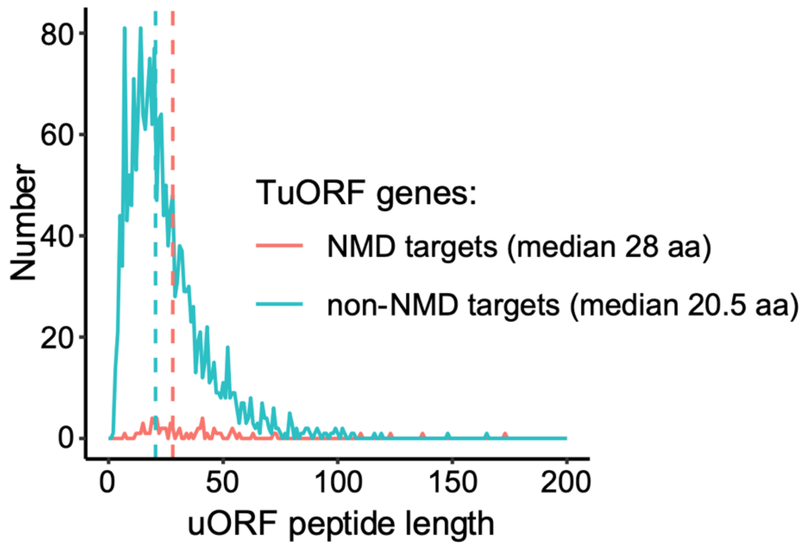


**Figure S8. Distributions of uORF peptide lengths in NMD target and non-NMD target TuORF genes.** Vertical dashed lines indicate the median value of each group.

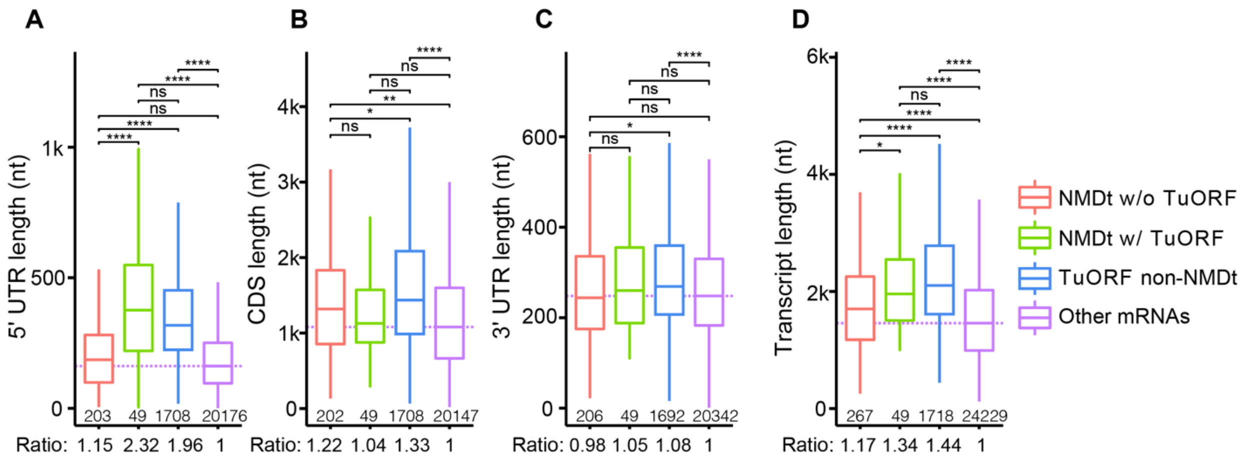


**Figure S9. Lengths of UTRs and CDSs for NMD targets and TuORF mRNAs.**
Lengths of the 5' UTRs (A), CDSs (B), 3' UTRs (C), and entire transcripts (D) of NMD targets (NMDt) without a TuORF, NMD targets with a TuORF, non-NMD TuORF mRNAs, and other mRNAs. Within the boxplots, the dashed lines mark the median level of other mRNAs; the number of genes in each category is listed below the lower whisker. The ratios underneath the plots indicate the median of each group normalized to that of other mRNAs. The statistical significance was determined as described in the Figure 2 legend.

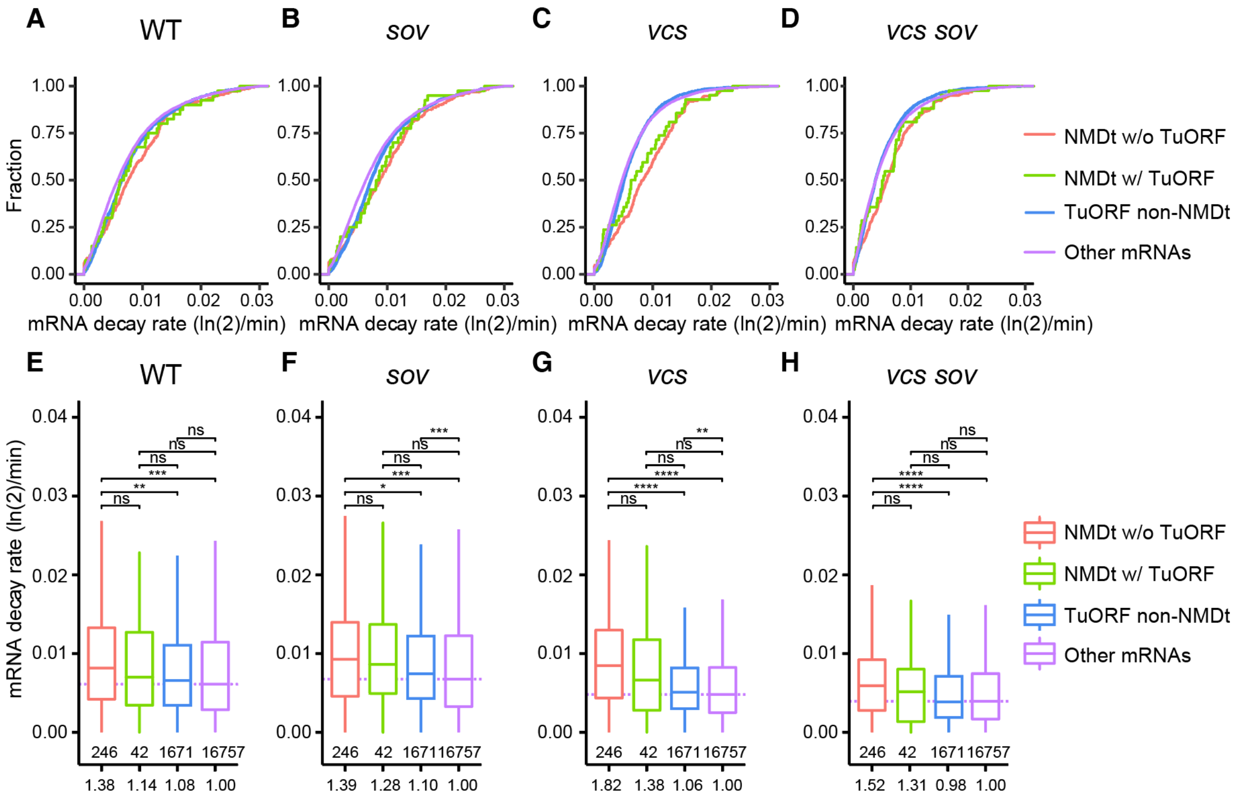


**Figure S10. The mRNA decay rates of NMD targets and TuORF-containing mRNAs in different genetic backgrounds.** Cumulative plots (A-D) and boxplots (E-H) showing the mRNA decay rates for NMD targets and TuORF mRNAs in wild type and mutants defective in either 5' to 3' decay (*vcs-7*), 3' to 5' decay (*sov*), or both (*vcs sov).* The mRNA decay rates were extracted from (Sorenson et al., 2018). Within the boxplots, the dashed lines mark the median level of other mRNAs; the number of genes in each category is listed below the lower whisker. The ratios underneath the plots indicate the median of each group normalized to that of other mRNAs. The statistical significance was determined as described in the Figure 2 legend.

**Reference:**

**Hsu, P.Y., Calviello, L., Wu, H.-Y.L., Li, F.-W., Rothfels, C.J., Ohler, U., and Benfey, P.N.** (2016). Super-resolution ribosome profiling reveals unannotated translation events in Arabidopsis. Proceedings of the National Academy of Sciences of the United States of America **113**: E7126–E7135.

**Papatheodorou, I. et al.** (2020). Expression Atlas update: From tissues to single cells. Nucleic Acids Research **48**: D77–D83.

**Song, G., Hsu, P.Y., and Walley, J.W.** (2018). Assessment and Refinement of Sample Preparation Methods for Deep and Quantitative Plant Proteome Profiling. Proteomics **18**: e1800220.

**Sorenson, R.S., Deshotel, M.J., Johnson, K., Adler, F.R., and Sieburth, L.E.** (2018). Arabidopsis mRNA decay landscape arises from specialized RNA decay substrates, decapping-mediated feedback, and redundancy. Proceedings of the National Academy of Sciences of the United States of America **115**: E1485–E1494.

**Wu, H.-Y.L., Song, G., Walley, J.W., and Hsu, P.Y.** (2019). Translational landscape in tomato revealed by transcriptome assembly and ribosome profiling. bioRxiv **181**: 367–380.
